# AAV-mediated interneuron-specific gene replacement for Dravet syndrome

**DOI:** 10.1101/2023.12.15.571820

**Authors:** John K. Mich, Jiyun Ryu, Aguan D. Wei, Bryan B. Gore, Rong Guo, Angela M. Bard, Refugio A. Martinez, Yemeserach Bishaw, Em Luber, Luiz M. Oliveira Santos, Nicole Miranda, Jan-Marino Ramirez, Jonathan T. Ting, Ed S. Lein, Boaz P. Levi, Franck K. Kalume

## Abstract

Dravet syndrome (DS) is a devastating developmental epileptic encephalopathy marked by treatment-resistant seizures, developmental delay, intellectual disability, motor deficits, and a 10-20% rate of premature death. Most DS patients harbor loss-of-function mutations in one copy of *SCN1A*, which has been associated with inhibitory neuron dysfunction. Here we developed an interneuron-targeting AAV human *SCN1A* gene replacement therapy using cell class-specific enhancers. We generated a split-intein fusion form of *SCN1A* to circumvent AAV packaging limitations and deliver *SCN1A* via a dual vector approach using cell class-specific enhancers. These constructs produced full-length Na_V_1.1 protein and functional sodium channels in HEK293 cells and in brain cells *in vivo*. After packaging these vectors into enhancer-AAVs and administering to mice, immunohistochemical analyses showed telencephalic GABAergic interneuron-specific and dose-dependent transgene biodistribution. These vectors conferred strong dose-dependent protection against postnatal mortality and seizures in two DS mouse models carrying independent loss-of-function alleles of *Scn1a,* at two independent research sites, supporting the robustness of this approach. No mortality or toxicity was observed in wild-type mice injected with single vectors expressing either the N-terminal or C-terminal halves of *SCN1A*, or the dual vector system targeting interneurons. In contrast, nonselective neuronal targeting of *SCN1A* conferred less rescue against mortality and presented substantial preweaning lethality. These findings demonstrate proof-of-concept that interneuron-specific AAV-mediated *SCN1A* gene replacement is sufficient for significant rescue in DS mouse models and suggest it could be an effective therapeutic approach for patients with DS.

## Introduction

Dravet syndrome (DS) is a severe early-onset epileptic encephalopathy marked by spontaneous and febrile seizures, motor disabilities, cognitive dysfunction, developmental delay, and heightened risk of premature death by sudden unexpected death in epilepsy (SUDEP)^1–3^. DS afflicts approximately 1:16000 births, usually manifests in the first year of life, and produces profound symptoms that require life-long care^4^. Most first-line anti-epileptic drugs are ineffective or contraindicated for DS^5^, although several recently approved drugs now partially ameliorate DS symptoms^6–8^. Critically, no FDA-approved long-term disease-modifying treatments currently exist for DS despite extensive efforts^9–11^. As a result, a treatment for DS is a pressing unmet need for patients and their caregivers.

Several aspects of DS pathophysiology have become clear. Over 80% of patients harbor monoallelic loss-of-function mutations in *SCN1A*^2,12–15^, which encodes Na_V_1.1, one of the voltage-gated sodium channels expressed in brain^16–18^. Mouse models with monoallelic *Scn1a* disruptions recapitulate the major clinical presentations of DS^19–24^, confirming genetic causality in DS. These mouse models suggest DS is associated with loss of excitability in telencephalic fast-spiking interneurons^19,25^, consistent with inhibition stimulation studies in patients^26^. Furthermore, targeted monoallelic *Scn1a* disruption in interneurons is sufficient for epileptic symptomology^24,27,28^, which can be ameliorated by simultaneous disruption in excitatory neurons^27^. Overall, these results suggest interneurons are the critical pathological cell population in DS and represent a putative effective target for cell class-specific therapies for this condition. Enhancer-adeno-associated viruses (AAVs)^29–33^ constitute a new technology that would enable interneuron-specific expression of a DS therapeutic transgene; however, AAVs are limited by their genome size constraints (∼4.7kb), which precludes delivery of human *SCN1A* open reading frame (ORF, 6.0kb) in a single AAV. Thus, AAV-mediated *SCN1A* gene replacement for DS has not been possible.

In this study, we demonstrate functional rescue in DS mice by restoring *SCN1A* to telencephalic GABAergic interneurons using AAV vectors. We achieved this by splitting the *SCN1A* ORF into two fragments (two “halves”) that undergo intein-mediated ligation to reconstitute a scarless, full-length functional voltage-gated sodium channel Na_V_1.1. With this split-intein mechanism and class-specific enhancers, we demonstrate interneuron-specific delivery and reconstitution of Na_V_1.1, which completely rescues mortality in DS model mice and confers strong resistance to epileptic seizures. Importantly, we show that expression of the transgenes or individual *SCN1A* halves in WT mice does not cause any overt toxicity. In contrast, we also find that nonselective neuronal expression of *SCN1A* generates early lethal toxicity. Together these data suggest telencephalic GABAergic interneuron-specific expression of *SCN1A* could provide a safe and effective therapeutic for DS.

## Results

### Split-intein fusion constructs produce full-length functional Na_V_1.1 channels

The open reading frame for human *SCN1A* (6.0 kb) is larger than the packaging limit of recombinant AAVs (∼4.7 kb^34^); this has thus far prevented an AAV gene replacement therapy for Dravet syndrome. To overcome this challenge, we divided the gene into two halves and used split-intein protein splicing to reconstitute the gene product (Na_V_1.1 channel) after translation^35,36^. We designed and built DNA constructs using this approach for a dual vector gene replacement therapy for Dravet syndrome in mice.

We developed split-intein DNA constructs using the predominantly expressed 1998-amino acid SCN1A isoform (**Fig. S1**) sequence which was codon optimized to maximize expression. We placed the split-intein breakpoint directly upstream of Cys1050 since intein-mediated protein ligation requires the presence of a cysteine residue adjacent to the split site^37^. We utilized the Cfa split-intein which was engineered for rapid activity and chemical stability^36^. The N- and C-terminals of the Cfa split-intein were respectively fused to the N- and C-terminals of halves of the split SCN1A gene (**Fig. 1A**). This approach leads to a scarless protein junction with no mutations at the splicing site. It has been successfully used previously to segmentally assemble a voltage-gated calcium channel^35^ which is structurally homologous to voltage-gated sodium channels. To the recombinant split-intein *SCN1A* halves, we added optimized short intron sequences within each coding sequence for improved expression, as well as HA and FLAG epitope tags for immunodetection (**Fig. 1A,B**).

**Figure 1:**
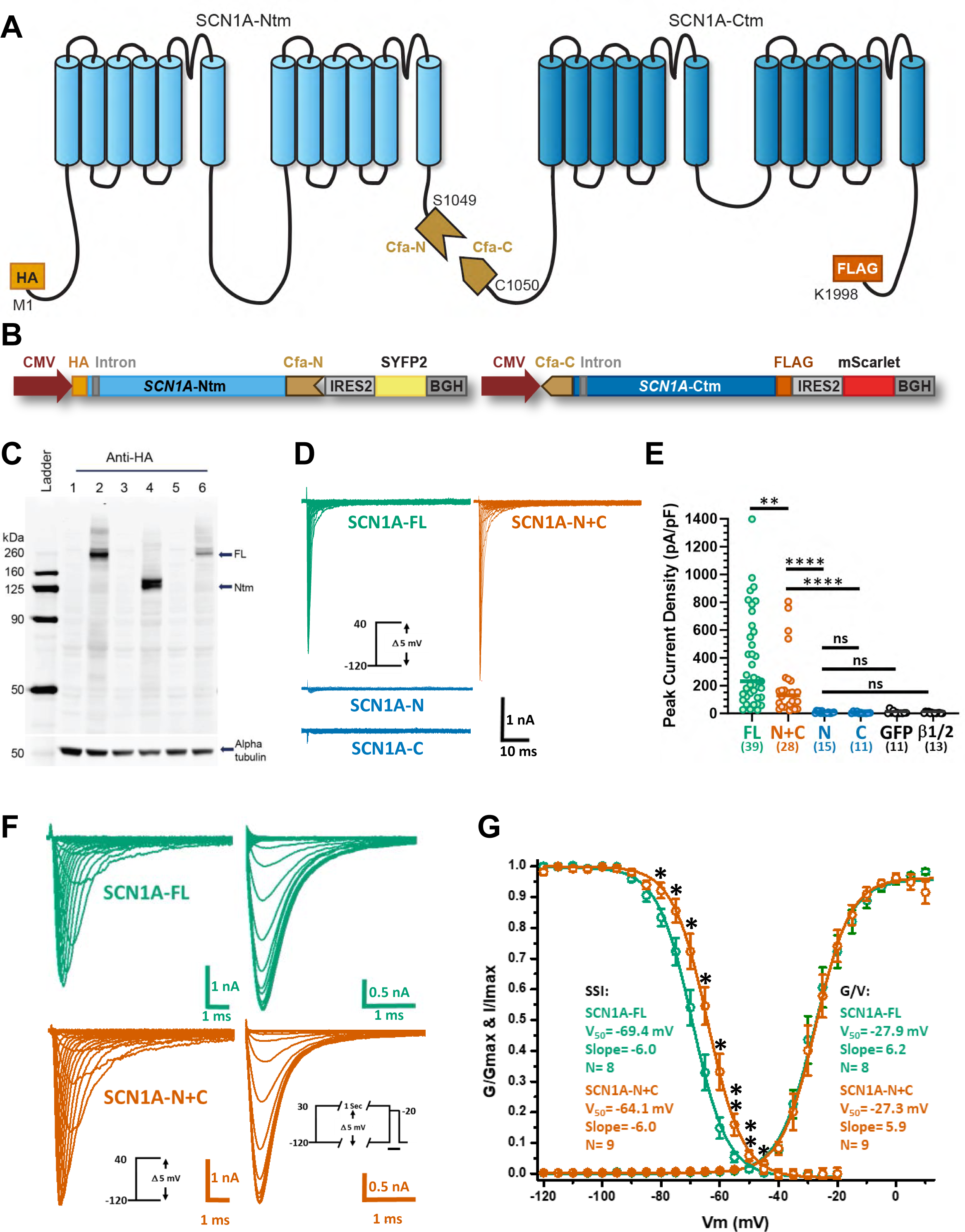
A functional split-intein design of SCN1A that reconstitutes functional Na_V_1.1 activity. (A) Design of split-intein fusion protein halves of SCN1A. We inserted the breakpoint and Cfa-N and Cfa-C split-intein peptides just before the native Cys1050 with no additional amino acids. After joining, the Cfa-N and Cfa-C intein fragments self-excise and yield scarless reconstituted SCN1A. HA and FLAG epitopes are inserted at the N- and C-termini of the N- and C-terminal halves for detection. (B) Cloning SCN1A split-intein fusion protein halves into CMV-driven plasmid vectors for testing functionality in cell lines, as well as IRES2-SYFP2 and IRES2-mScarlet transfection reporters. (C) Reconstitution of full-length SCN1A after co-transfection of split-intein fusion protein halves into HEK-293 cells. We analyzed whole cell protein preparations by western blot for HA epitope tag after transfection. Lanes: 1) empty vector, 2) full-length HA-SCN1A, 3) full-length SCN1A-FLAG, 4) HA-SCN1A-Ntm, 5) SCN1A-FLAG-Ctm, and 6) HA-SCN1A-Ntm plus SCN1A-FLAG-Ctm. Expected sizes: full-length HA-SCN1A 232 kDa, HA-SCN1A-Ntm 134 kDa, reconstituted full-length HA-SCN1A-FLAG after joining 233 kDa. Anti-tubulin is a loading control. (D) Exemplary currents evoked in HEK293 cells transfected with full-length SCN1A (SCN1A-FL, green), a combination of SCN1A-Ntm and SCN1A-Ctm (SCN1A-N+C, orange), SCN1A-Ntm only (SCN1A-N, blue) and SCN1A-Ctm only (SCN1A-C, blue), in response to a family of step depolarizations from a holding potential of -120 mV to 40 mV, in 5 mV increments. Co-transfection with SCN1A-Ntm and SCN1A-Ctm produced functional Na_V_1.1 currents comparable in size to full-length SCN1A. Scale bar: 1 nA, 10 msec. (E) Peak current densities, normalized to capacitance, for full-length SCN1A (FL, n= 39), SCN1A-Ntm + SCN1A-Ctm (N+C, n= 28), SCN1A-Ntm (N, n= 15), SCN1A-Ctm (C, n= 11), GFP/empty vector (GFP, n= 11) and SCN1B/2B (β1/2, n= 13). Medians displayed with data points. FL and N+C datasets are significantly different (**, p= 0.0083). N+C is highly significantly different compared to N or C (****, p< 0.0001). N and C are not significantly different from GFP and β1/2 controls (p> 0.05, ns). Statistical comparisons performed by pairwise Mann-Whitney U tests. All HEK293 transfections performed in the background of a separate SCN1B/2B expression plasmid, at a ratio of 10:1. (F) Voltage-dependent gating properties of WT full-length human SCN1A (SCN1A-FL) and reconstituted SCN1A channels formed by co-expression of N-terminal SCN1A-Cfa intein (SCN1A-N) and Cfa intein-C-terminal SCN1A (SCN1A-C) plasmid constructs, acutely expressed in HEK293 cells and characterized by whole-cell patch-clamp recordings. Currents activated by a family of step depolarizations from SCN1A-FL (green) and reconstituted SCN1A-N+C (orange), from a holding potential of -120 mV (left panels). Steady-state inactivation current traces evoked by a step to -20 mV, after a family of 1 sec preconditioning steps from -120 mV to 30 mV (right panels). Scale bars for activation currents (F, left panels) equal 1.0 nA, and for SSI currents (F, right panels) equal 0.5 nA; time equals 1.0 msec for all panels. (G) Conductance-voltage (G/V) and steady-state inactivation (SSI) plots for SCN1A-FL (green) and reconstituted SCN1A-N+C (orange) channels, fitted to single Boltzmann functions. G/V plots for both constructs are indistinguishable, whereas the SSI plot for SCN1A-N+C is shifted by +5 mV relative to SCN1A-FL. Statistical significance derived from unpaired pairwise t-tests, assuming equal variance (* p< 0.05; ** p< 0.01).

To confirm intein-mediated reconstitution of full-length Na_V_1.1 channels, the *SCN1A* gene halves were expressed in HEK-293 cells using the CMV promoter (**Fig. 1B,C**). By western blot analysis, cells transfected with HA-tagged full-length *SCN1A* produced a single band with an apparent mass of approximately 250 kDa (predicted mass 232 kDa, **Fig. 1C** Lane 2). Those transfected with HA-tagged SCN1A-Ntm half led to an apparent mass of approximately 135 kDa (predicted mass 134 kDa, **Fig. 1C** Lane 4). However, cells transfected with both SCN1A-Ntm and SCN1A-Ctm halves demonstrated a strong band at approximately 250 kDa and only a weak band at approximately 135 kDa (**Fig. 1C** Lane 6). These results demonstrate that our constructs led to efficient intein-directed ligation of Na_V_1.1 protein halves, similar to other reported split-protein constructs using the same Cfa split intein^36,38^.

To examine whether reconstituted Na_V_1.1 protein derived from split-intein *SCN1A* halves was functional, we assessed HEK-293 cells expressing these constructs by whole-cell patch clamp electrophysiology. Fluorescent protein reporters appended to each *SCN1A* expression construct were used to identify transfected cells and confirm expression (**Fig. 1B**). To promote Na_V_1.1 channel cell surface expression, constructs were co-transfected with Na_V_1.1 β-subunits (SCN1B/SCN2B^16^, see methods). We observed that cells expressing full length *SCN1A* (SCN1A-FL) constructs show rapid and large inward sodium currents in response to depolarizing voltage steps as previously demonstrated^16,18,39^ (**Fig. 1D**). Cells expressing both *SCN1A-Ntm* and *SCN1A-Ctm* (SCN1A-N+C) also showed large inward sodium currents, but not cells expressing either halves alone or empty vector (**Fig. 1D**). Quantification revealed significantly greater capacitance-normalized peak current densities in SCN1A-N+C cells (median current density 130.6 pA/pF) than cells singly expressing either *SCN1A-Ntm* or *SCN1A-Ctm* alone or negative controls (SCN1A-N 4.5 pA/pF, p< 0.0001; SCN1A-C 3.3 pA/pF, p< 0.0001; GFP/empty vector 3.7 pA/pF, p= 0.0084; β-subunits alone 2.3 pA/pF, p= 0.0038, all pairwise Mann-Whitney U tests, **Fig. 1E**). Median peak current density for SCN1A-N+C cells (130.6 pA/pF) was significantly less than that observed in SCN1A-FL cells (232.0 pA/pF) (p= 0.0083, pairwise Mann-Whitney U test), possibly due to less of each DNA construct used for transfections (0.4 μg each of *SCN1A-Ntm* and *SCN1A-Ctm*, as versus 0.8 μg for SCN1A-FL). Examinations of the records at faster time scales show that both SCN1A-FL and SCN1A-N+C cells expressed similarly rapidly activating and inactivating inward currents in response to step depolarizations (**Fig. 1F**).

To quantify whether the current mediated by split-intein reconstructed channels exhibit the known normal voltage-dependent gating properties of Na_V_1.1 channels, we analyzed conductance-voltage (G/V) activation and steady-state inactivation (SSI) relationships. Indistinguishable G/V activation plots were exhibited by cells transfected with SCN1A-FL (V_50_= - 27.9 mV, slope= 6.2) or SCN1A-N+C (V_50_= -27.3 mV, slope= 5.9), similar to previously reported full-length *SCN1A*^16^ (V_50_= -26.4 mV, slope= 7.1). Steady-state inactivation profiles for SCN1A-FL and SCN1A-N+C also were both similar to that previously reported for full-length *SCN1A* (V_50_= -67.5 mV, slope= -6.2)^16^. The SSI profile for reconstituted SCN1A-N+C currents (V_50_= - 64.1 mV, slope= -6.0) was observed to be slightly but significantly depolarized by approximately 5 mV relative to SCN1A-FL (V_50_= -69.4 mV, slope= -6.0) (**Fig. 1G**). Overall, these results demonstrate that functional Na_V_1.1 sodium channels are efficiently reconstituted from our two half *SCN1A* split-intein constructs, with similar voltage-dependent gating properties to full-length *SCN1A*, and this enables delivery of Na_V_1.1 channel activity from two AAV-sized vectors.

### DLX2.0-driven intein-SCN1A vectors specifically transduce telencephalic GABAergic interneurons

We cloned the split-intein fusion protein halves HA-SCN1A-Ntm and SCN1A-Ctm-FLAG into AAV plasmid vectors under control of an optimized hDLXI5/6i enhancer^29,40^ (“DLX2.0”) that drives transgene expression in telencephalic GABAergic interneurons in mice and in human organotypic slice cultures^31,41^. These constructs were packaged into research-grade AAV2/PHP.eB viral vectors^42^ at Packgene Inc. (Guangzhou, China, and Houston, TX, USA; **Fig. 2A**). Neonatal mice were injected at postnatal day 2 (P2) with 5 μL total volume of these vectors delivered bilaterally via intracerebroventricular (ICV) route at a dose of 3e10 genome copies (gc) of each half. After twenty days, we analyzed mouse motor cortex membrane protein content by western blot to assess efficiency of half joining by intein-mediated fusion. Mice injected with either DLX2.0-HA-SCN1A-Ntm or DLX2.0-SCN1A-Ctm-FLAG PHP.eB AAVs alone showed weak HA- or FLAG-immunoreactive bands near their expected sizes of their protein products (N-term predicted size 134 kDa, C-term predicted size 115 kDa, **Fig. 2B**). The weakness of the unjoined SCN1A half-protein bands in our membrane protein fraction is likely a technical artifact of the western blot membrane protein preparation since half proteins alone are efficiently detected by immunohistochemistry (IHC) with no obvious change in subcellular distribution when co-delivered (see **Fig. 2C** below). Regardless, when we co-injected viral vectors for both halves, we observed strong bands near the expected apparent size of intact Na_V_1.1 near 250 kDa, demonstrating that intein-mediated reconstitution of full-length human Na_V_1.1 occurs in mouse telencephalic GABAergic interneurons (**Fig. 2B**).

**Figure 2:**
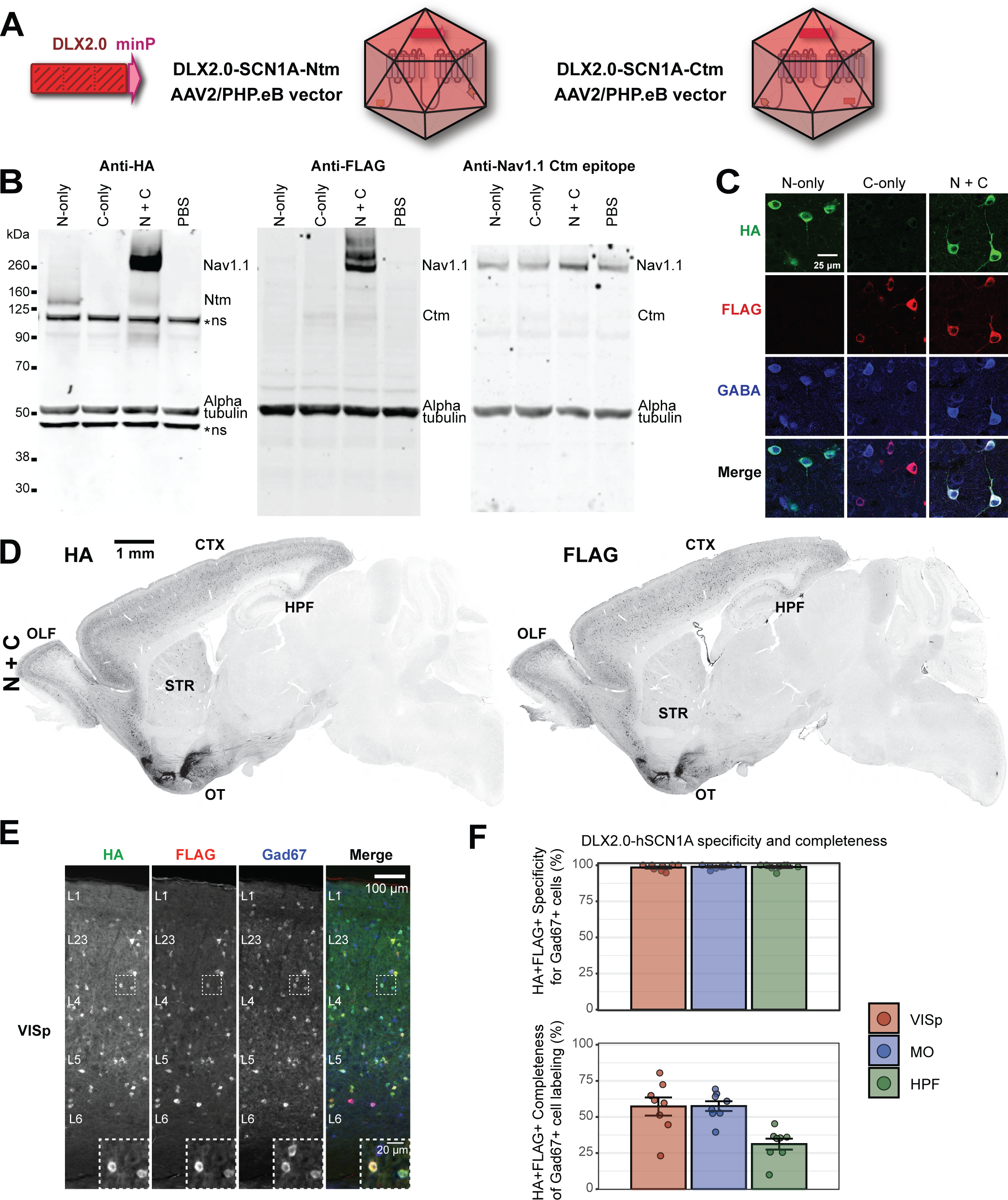
Cell class-specific delivery of SCN1A to telencephalic GABAergic interneurons using optimized enhancer DLX2.0. (A) Recombinant AAV2/PHP.eB vectors for delivery of DLX2.0-split-intein-SCN1A. (B) Efficient SCN1A reconstitution in mouse brain with DLX2.0-split-intein-SCN1A vectors. In panels B-F we injected P2 neonatal mice BL-ICV with 3e10 gc of each indicated vector. At P24 we analyzed mouse brain membrane protein fractions by western blotting with antibodies targeting HA or FLAG epitope tags, the C-terminus of Na_V_1.1, and alpha-tubulin as a loading control. Note the PBS-injected negative control is the same lane as that shown in Fig. 6B as these experiments were performed together. (C) Specific detection of HA- and FLAG-expressing GABA+ cells in cortex. HA- and FLAG-expressing cells can be observed when either half is delivered alone or together. Layer 2/3 of VISp is shown at P20. (D) Representative stitched fluorescence image of biodistribution of HA and FLAG epitopes in scattered telencephalic neurons. Expression is pseudo-colored black. (E) Representative stitched fluorescence image of HA and FLAG epitopes, and Gad67+ neurons in VISp. In panels D-E we show expression at P47. (F) High specificity and completeness of expression within Gad67+ neurons in multiple telencephalic regions across multiple animals. We counted cells that express both HA and FLAG epitopes in visual VISp, MO, and HPF. Layer 1 was excluded from VISp and MO analysis due to DLX2.0-PHP.eB vectors inefficiently targeting that layer^31^. Each point represents one mouse, bars represent the means, and error bars represent standard error of the mean. Mice span ages P47-P139, mean age P85. Abbreviations: CTX cerebral cortex, OLF olfactory areas, HPF hippocampal formation, OT olfactory tubercle, STR striatum, VISp primary visual cortex, MO motor cortex.

We also analyzed the biodistribution of transgene expression by IHC in animals injected with these vectors. We observed strong HA and FLAG immunoreactivity in scattered telencephalic neurons of mice injected with each vector alone, or when injected together (**Fig. 2C**). Both HA- and FLAG-expressing cells were present throughout the telencephalon after BL-ICV delivery (**Fig. 2D**). Co-staining with the GABAergic neuron markers GABA and Gad67 demonstrated high overlap of HA- and FLAG-expressing cells (**Fig. 2C, E**). Quantitative analysis of these IHC data showed that 98-99% of cells co-expressing HA and FLAG were Gad67+ in several telencephalic regions (**Fig. 2F**), indicating high specificity as seen in other studies using the DLX2.0 enhancer^31,41^. Additionally, HA+FLAG+ expression was observed in a substantial proportion of the telencephalic Gad67+ GABAergic neuron population in different brain regions, including those known to be involved in seizure generation such as hippocampus and cortex (mean ± standard deviation: VISp 57 ± 18%, MO 57 ± 10%, HPF 31 ± 11%, n= 8 mice, **Fig. 2F**). Using a separate cohort of *Dlx5/6-Cre; Ai14* mice to provide an independent label for telencephalic GABAergic interneurons^43,44^, we observed similar high levels of specificity and moderate completeness of AAV transduction (**Fig. S2**). These data demonstrate that our vectors effectively deliver both the C- and N-terminal split-intein *SCN1A* halves with high specificity and moderate coverage for telencephalic GABAergic interneurons in mice. Together with our western blot and electrophysiology results above, these findings indicate that upon their expression, the two protein products of *SCN1A* halves fuse using the split-intein leading to expression of functional full-length Na_V_1.1 channels in telencephalic GABAergic interneurons.

### Dual DLX2.0 split-intein SCN1A vectors protect against SUDEP in DS mouse models

With the ability to deliver SCN1A with cell class specificity, we next tested whether telencephalic GABAergic interneuron-specific SCN1A gene replacement could rescue DS symptoms in mouse models. We used *Scn1a^fl/+^;Meox2-Cre DS* mice bred on a pure C57Bl/6 background^24,45,46^ housed at Seattle Children’s Research Institute. These animals demonstrate ∼50% mortality due to SUDEP by P70^45^, similar to other DS model lines^20,47–49^, analogous to the ∼15% rate of SUDEP in DS patients^3^. We injected a cohort of neonatal DS model mice (BL-ICV at P0-P3 with 3e10 gc each vector of DLX2.0-split-intein-SCN1A produced at PackGene Inc.) and measured mortality and susceptibility to thermally induced seizures in these animals (**Fig. 3A**). Remarkably, all treated DS mice (n= 27/27) survived beyond P70, as opposed to negative control mice either untreated or receiving control AAVs (empty or single part-only vectors), which showed significantly higher mortality (Untreated n= 34/68, 50% mortality, Log-rank test p= 1.4e-5; Empty/single-part n= 15/40, 37.5% mortality, Log-rank test p= 6.6e-4; **Fig. 3B**). No side effects were observed in littermate control mice that received dual vector treatment. Strikingly, a subset of the treated DS animals was maintained longer, and 100% survived to beyond P200 (n= 11/11). These findings demonstrate that telencephalic GABAergic interneuron supplementation of AAV-mediated SCN1A transgene is sufficient to completely prevent SUDEP in DS model mice. *Dual DLX2.0 split-intein SCN1A vectors protect against thermal and spontaneous myoclonic and generalized tonic-clonic seizures in DS mice*. DS model mice are sensitive to thermally induced seizures, similar to patients with DS^1,2,19,46^. We previously showed that small elevations of core body temperature trigger several myoclonic (MC) seizures, leading ultimately to a generalized tonic-clonic (GTC) seizure in DS mice^46^. To examine the efficacy of our DLX2.0- SCN1A vectors in preventing thermally induced MC and GTC seizures, we analyzed DS mice between P25 to P35 (**Fig. 3A**). First, video analysis revealed that mice treated with dual DLX2.0-SCN1A AAVs were protected from MC seizures up to 40⁰C (n= 16/16), at which temperature fewer than half of negative control mice were MC seizure-free (Untreated n= 5/11, Empty/single part AAVs n= 9/23, **Fig. 3C**). With further increase in temperature, a growing fraction of the treated mice began to exhibit MC seizures, reaching 50% of treated subjects at 42⁰C (n= 8/16), but at this temperature significantly greater numbers of negative control mice experienced MC seizures (Untreated n= 9/11, Fisher’s exact test p= 0.042; Empty/single part AAVs n= 21/23, Fisher’s exact test p= 0.011, **Fig. 3C**). Second, we also observed treatment conferred GTC seizure protection in this assay. All treated DS mice were free from GTC seizures up to 41.5⁰C (n= 16/16, 100%, **Fig. 3D**), and the majority were protected from GTC seizures at 42⁰C, the last stage of the test (n= 15/16, 94%), which was significantly greater than negative control treatment groups (Untreated n= 4/11, Fisher’s exact test p= 0.0025; Empty/single part AAVs n= 7/23, Fisher’s exact test p= 0.00010, **Fig. 3D**). Finally, assessment of all the MC seizures preceding a GTC seizure showed that the cumulative MC event number before GTC seizure onset was significantly suppressed when DS mice were treated with dual DLX2.0-SCN1A AAVs compared to control AAV treatment or untreated controls (Untreated or Empty/single versus DLX2.0-SCN1A, unpaired t-test p< 0.01 at each temperature from 39.5 to 41, **Fig. 3E**). These findings demonstrate that DLX2.0 mediated delivery of SCN1A transgene specifically to telencephalic GABAergic interneurons yields substantial protection from thermally induced seizures.

**Figure 3:**
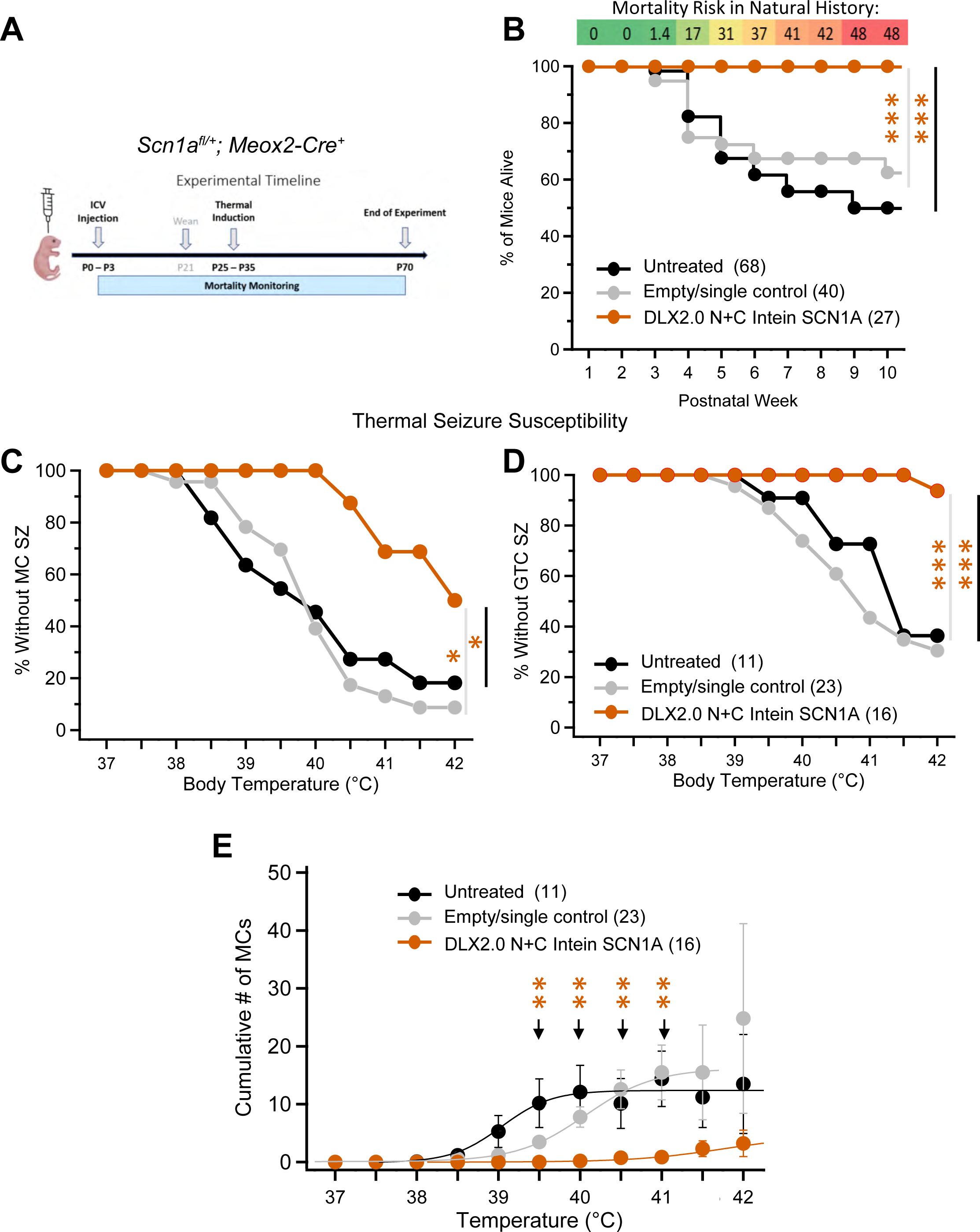
Recovery of mortality and epileptic symptoms in DS model mice with DLX2.0-split intein-SCN1A. (A) Experimental timeline to rescue of epileptic symptoms in DS model mice. We used *Scn1a^fl/+^; Meox2-Cre* animals on a pure C57BL/6 background to model DS^45^. We injected P0-P3 pups with empty control, single-part alone control, or telencephalic GABAergic interneuron-targeting dual DLX2.0 N+C SCN1A AAVs (3e10 gc each vector per animal), and monitored mortality until P70. Some animals were tested for seizure susceptibility by thermal challenge between P25-P35. (B) Mortality protection in DS model mice after treatment with DLX2.0-SCN1A AAVs. Mice treated with DLX2.0 N+C SCN1A AAVs (n= 27) exhibit significantly greater survival than untreated mice (n= 68, *** Log-rank test p= 1.4e-5) or mice treated with empty or single-part vector controls (n= 40, *** Log-rank test p= 6.6e-4). Note the untreated control groups in **Fig. 3B-E** represents the same set of untreated animals as that shown in Fig. 7B, D-F. (C-E) Protection from heat-induced seizures in DS model mice after treatment with DLX2.0-SCN1A AAVs. (C) DS model mice treated with DLX2.0 N+C SCN1A AAVs (n= 16) are significantly less likely to exhibit MC seizures by 42°C than untreated mice (n= 11, * Fisher’s exact test p= 0.042) or mice treated with empty or single-part vector controls (n= 27, * Fisher’s exact test p= 0.011). (D) DS model mice treated with DLX2.0 N+C SCN1A AAVs exhibit significantly less likely to exhibit GTC seizures by 42°C than untreated mice (** Fisher’s exact test p= 0.0025) or mice treated with empty or single-part vector controls (** Fisher’s exact test p= 0.00010). (E) DS model mice treated with DLX2.0 N+C SCN1A AAVs exhibit significantly fewer MC events during thermal challenge assay than untreated mice or mice treated with empty or single-part vector controls (** unpaired t-test p< 0.001 at each indicated timepoint, for comparison to Untreated and Empty/single part negative control animals).

To assess the effectiveness of DLX2.0-SCN1A AAV vectors to protect against spontaneous epileptic symptoms, we implanted untreated and treated DS model mice with electrodes for electrocorticography (ECoG) recordings. Untreated DS model mice displayed high-amplitude interictal spikes as seen in our previous work^45,50^ (**Fig. 4A**), and the frequency of these spikes was significantly diminished by dual DLX2.0-SCN1A AAV treatment (ANOVA p < 0.05, **Fig. 4B**). We also observed MC events during recordings of untreated DS model mice (n = 10/10, **Fig. 4C**), which were absent in recordings from treated DS model mice (n = 0/10, p = 7.1e-4 Fisher’s exact test, **Fig. 4D**). Finally, some untreated DS model mice exhibited spontaneous GTC seizures during recording (n = 2/10, **Fig. 4E**), but none of the treated animals exhibited GTCs (n = 0/10, **Fig. 4F**) although this effect was not significant (p = 0.47 Fisher’s exact test). These results indicate that dual DLX2.0-SCN1A AAV treatment can yield protection against spontaneous epileptic symptoms in DS model mice.

**Figure 4:**
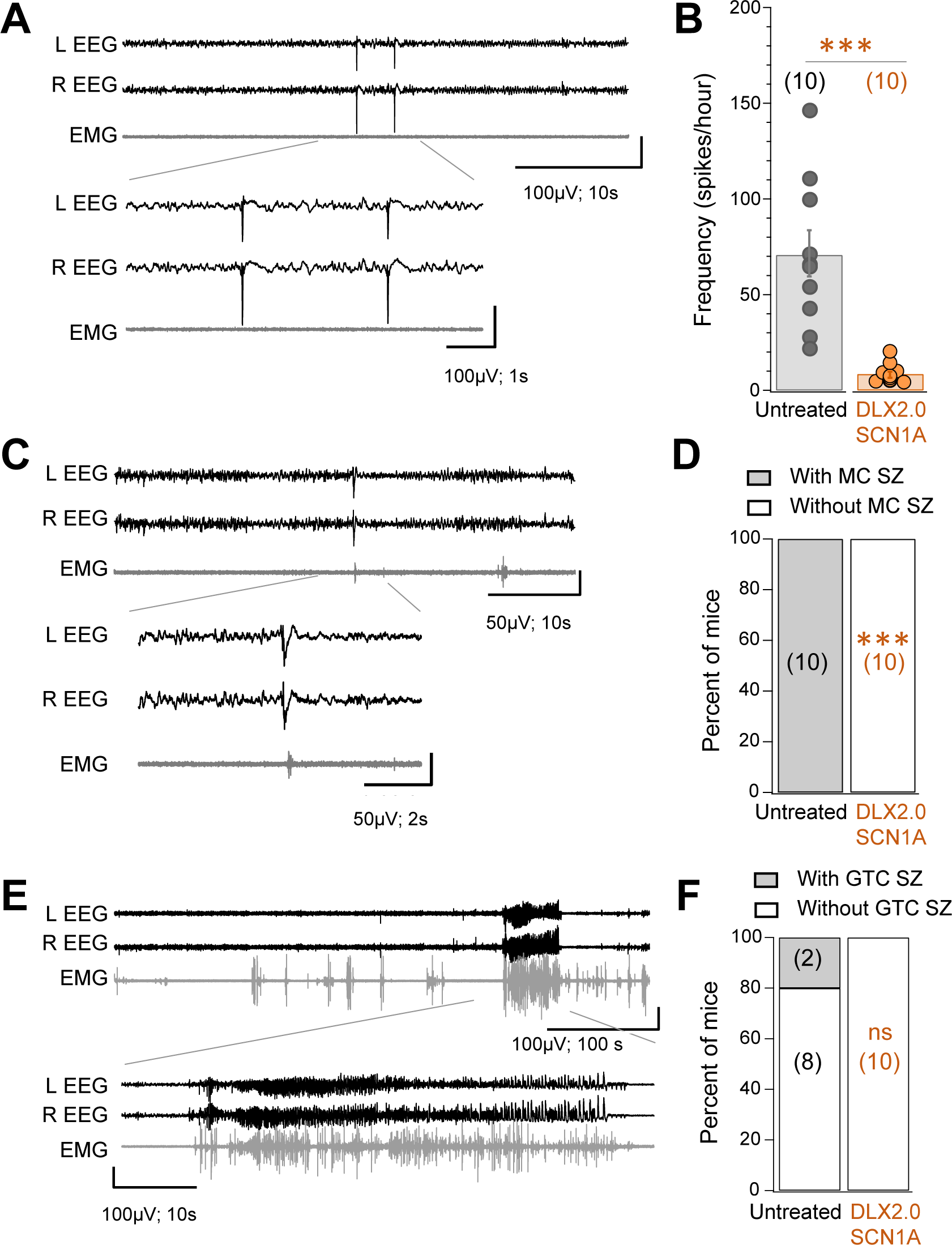
Recovery of spontaneous epileptic symptoms in DS model mice with DLX2.0-SCN1A AAVs. (A-B) Interictal spike reduction in DS model mice. We implanted *Scn1a^fl/+^; Meox2-Cre* DS model mice with ECoG/EMG electrodes, which revealed frequent interictal spikes generalized across the brain in both left (L) and right (R) channels as in our previous work^45,50^. (A) Example interictal spikes are shown in untreated mice. (B) Counting interictal spikes in untreated and treated (3e10 gc each DLX2.0 N+C SCN1A AAV vector delivered BL-ICV at P0-P3) mice reveals a significantly decreased frequency of interictal spikes with treatment. n= 10 animals per condition, each circle represents one animal, bars and error bars represent means and standard error of the means. *** p = 0.0001 by two-tailed unpaired t-test. (C-D) Spontaneous MC event prevention in DS model mice. (C) Example MC event observed in untreated *Scn1a^fl/+^; Meox2-Cre* DS model mice, which have sharp spikes followed closely by EMG signals. (D) We categorized mice as having MC events or not having MC events during the recording period, which revealed a highly significant prevention of MCs in the treated animals. *** p = 7.1e-4 by Fisher’s exact test. (E-F) Spontaneous GTCs in DS model mice. (E) Example spontaneous GTC seizure observed in an untreated *Scn1a^fl/+^; Meox2-Cre* DS model mice. (F) We categorized mice as having or not having GTC seizures during the recording period, which revealed treated animals did not exhibit GTC seizures although this effect was not significant (ns, p = 0.47 by Fisher’s exact test).

### Reproducibility across AAV batches

To confirm these findings, we tested independent batches of DLX2.0-split-intein-SCN1A vectors produced in-house at Allen Institute, and we observed dose-dependent specific expression in telencephalic GABAergic interneurons (**Fig. S3A-C**), which correlated with dose-dependent increases in survival (**Fig. S3D**) and thermal seizure protection (**Fig. S3E**). These results suggest reproducibility of epileptic rescue with independent batches of vector, although this in-house-packaged vector showed reduced levels of transduction and rescue as compared to the commercially produced vector.

### Reproducibility across mouse models and testing sites

To further confirm the therapeutic effects were robust, we retested the PackGene batch of DLX2.0-SCN1A AAV vectors in a second independent *Scn1a^+/R613X^*mouse model cohort on 129/BL6 F1 background housed at the Allen Institute for Brain Science (**Fig. S4A**). Consistent with previous reports^21,48^, we observed extensive premature lethality by P70 (n= 19/33, 58%) in uninjected mice, but 100% survival in mice injected with the dual DLX2.0-SCN1A AAVs (n= 30, p< 0.001 by Mantel-Cox test, **Fig. S4B**). After mortality monitoring, some mice underwent ECoG recordings which revealed untreated DS model mice demonstrate spontaneous GTC seizures and interictal spikes between P70 to P120 (**Fig. S4C**), which were both significantly reduced by the administration of DLX2.0-SCN1A vectors (GTCs, Mann-Whitney U test p= 0.020; spikes, Mann-Whitney U test p= 0.033; **Fig. S4C-E**). A separate subset of this cohort was monitored for long-term survival, and the dual-vector injected DS model mice exhibit 100% survival to beyond P365 (n= 14/14), with no evidence of toxicity in littermate control animals receiving either or both halves of the DLX2.0-SCN1A dual AAV vector system (**Fig. S4B**). Thus, DLX2.0-SCN1A AAV vectors conferred strong and reproducible protection from mortality, induced seizures, and spontaneous seizure burden in two independent genetic mouse models of DS, although this protective effect of the AAV treatment is dose- and AAV quality-dependent.

### Dual DLX2.0 vectors completely protect against SUDEP and seizures in mice with telencephalic GABAergic interneuron-specific Scn1a deletion

Previous studies have demonstrated much more severe mortality with specific *Scn1a* loss in interneurons, likely due to a disrupted excitatory/inhibitory balance^24,27,28^. To further investigate the effectiveness of the dual vector DLX2.0-SCN1A AAV, we delivered the AAVs to mice carrying the disease-causing mutation in the same telencephalic GABAergic interneuron population. These mice were generated by crossing mice carrying the *Dlx5/6-Cre* allele^43^ with mice carrying floxed *Scn1a*^24^ (**Fig. 5A**). Since the site of disease pathogenesis precisely matches the treatment target, in this experiment we directly tested the therapeutic effectiveness of the DLX2.0-SCN1A AAV vectors in a more precise rescue scenario. *Scn1a^fl/+^; Dlx5/6-Cre* mice were injected with dual DLX2.0-SCN1A AAVs via BL-ICV route at P0-3 and monitored these and untreated controls for premature death up to P70. Untreated mice showed severe mortality starting at postnatal week 3, with all the mice succumbing by week 6 (n= 31/31, **Fig. 5B**). In striking contrast, all mice treated with dual DLX2.0-SCN1A AAVs survived up to P70 (n= 9/9, p= 3.3e-14, Fisher’s exact test, **Fig. 5B**), despite the more severe adverse phenotype compared to mice with a global heterozygous loss of *Scn1a*. Furthermore, none of the treated mice exhibited either MC or GTC seizures during thermal challenge up to 42⁰C, unlike untreated animals (MCs: n= 8/10 untreated versus 0/9 treated, p= 7.1e-4; GTCs n= 10/10 untreated versus 0/9 treated, p= 1.1e-5; both Fisher’s exact test; **Fig. 5C**). These data demonstrate that rescue of DS phenotypes is possible when the therapeutic transgene is precisely delivered to the critical cell populations carrying disease-causing mutations, even in the face of more severe symptoms.

**Figure 5:**
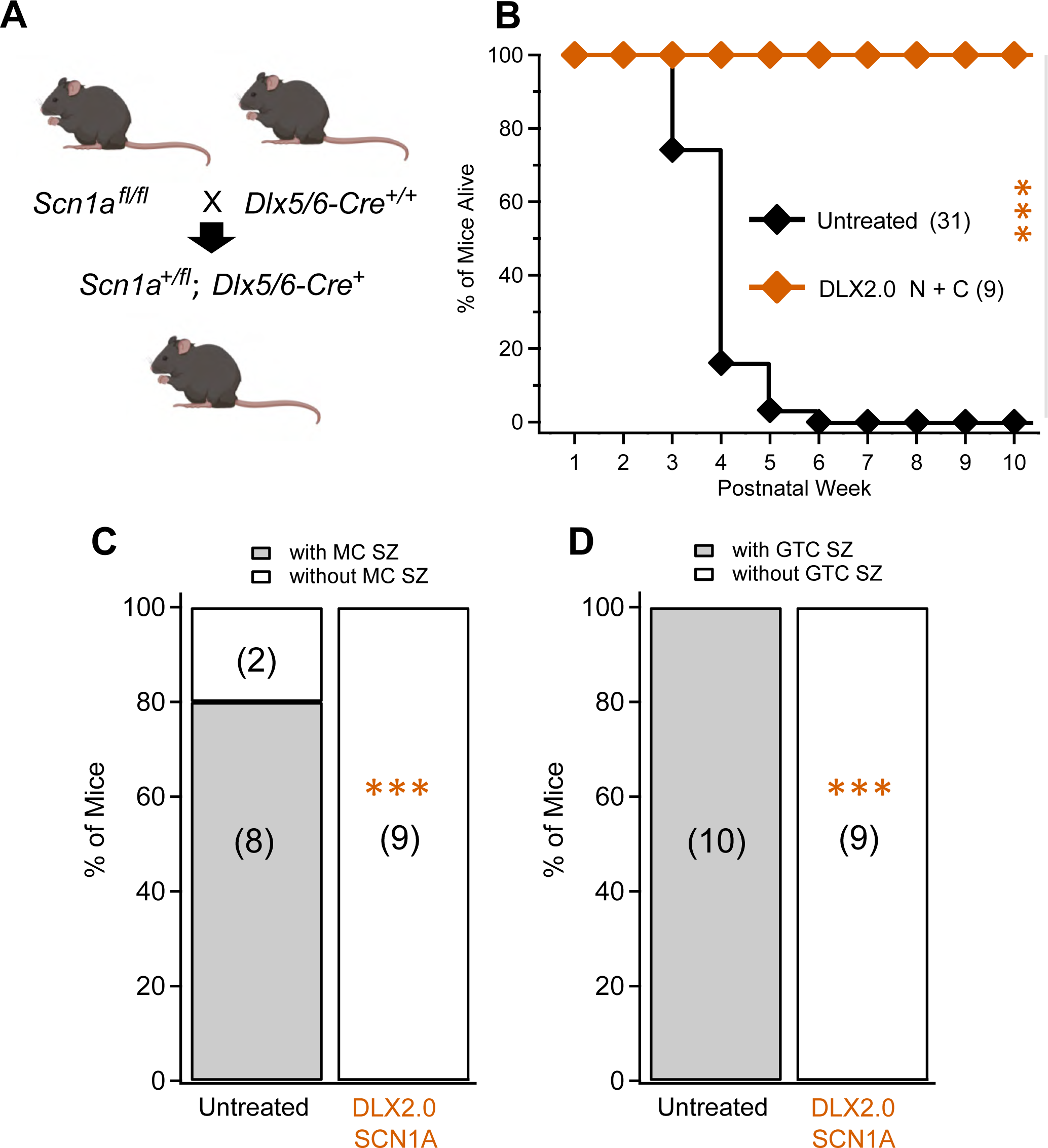
Recovery of severe epileptic phenotypes in mice lacking SCN1A in telencephalic GABAergic interneurons using DLX2.0-SCN1A AAVs. (A) Genetic cross resulting in 100% *Scn1a^+/fl^; Dlx5/6-Cre* animals. (B) Survival curves for treated and untreated *Scn1a^+/fl^; Dlx5/6-Cre* animals. Untreated pups exhibit 100% mortality by the 6^th^ week of life (n= 31/31). In contrast, pups injected with DLX2.0-split-intein-SCN1A (3e10 gc each vector at P2 by BL-ICV) exhibit 100% survival through the 10^th^ week of life (n= 9/9). *** p= 3.3e-14 by Fisher’s exact test. (C-D) Protection from seizures in treated *Scn1a^+/fl^; Dlx5/6-Cre* animals during thermal challenge assay. (C) Significant protection from thermally induced MC seizures (*** p= 7.1e-4, Fisher’s exact test). (D) Significant protection from thermally induced GTC seizures (*** p= 1.1e-5, Fisher’s exact test).

### hSyn1-driven vectors lead to nonselective neuronal expression of SCN1A

To determine whether telencephalic GABAergic interneuron-selective targeting is beneficial for gene replacement therapy in DS, as a comparator we produced and tested split-intein vectors driven by hSyn1 which expresses in most brain neuronal populations^51^, including excitatory and inhibitory neurons (**Fig. 6A**). Constructs for these vectors were built and packaged in the same way as DLX2.0 ones, except that hSyn1 promoter was used in this case. Delivered by BL-ICV at P2, these hSyn1 split-intein vectors led to reconstitution of full-length Na_V_1.1 in mouse brain (**Fig. 6B**), with expression observed in both Gad67+NeuN+ and Gad67-NeuN+ neurons (**Fig. 6C**). This expression pattern was observed throughout the telencephalon, with little expression seen in subtelencephalic structures likely due to the forebrain-biased delivery route (**Fig. 6D**). Quantification confirmed all labeled cells to be NeuN+ neurons (**Fig. 6E-F**). Average levels of completeness for HA+FLAG+ cells ranged from 6-18% NeuN+ cells and 13-33% Gad67+ cells, depending on the telencephalic structure analyzed (**Fig. 6F**).

**Figure 6:**
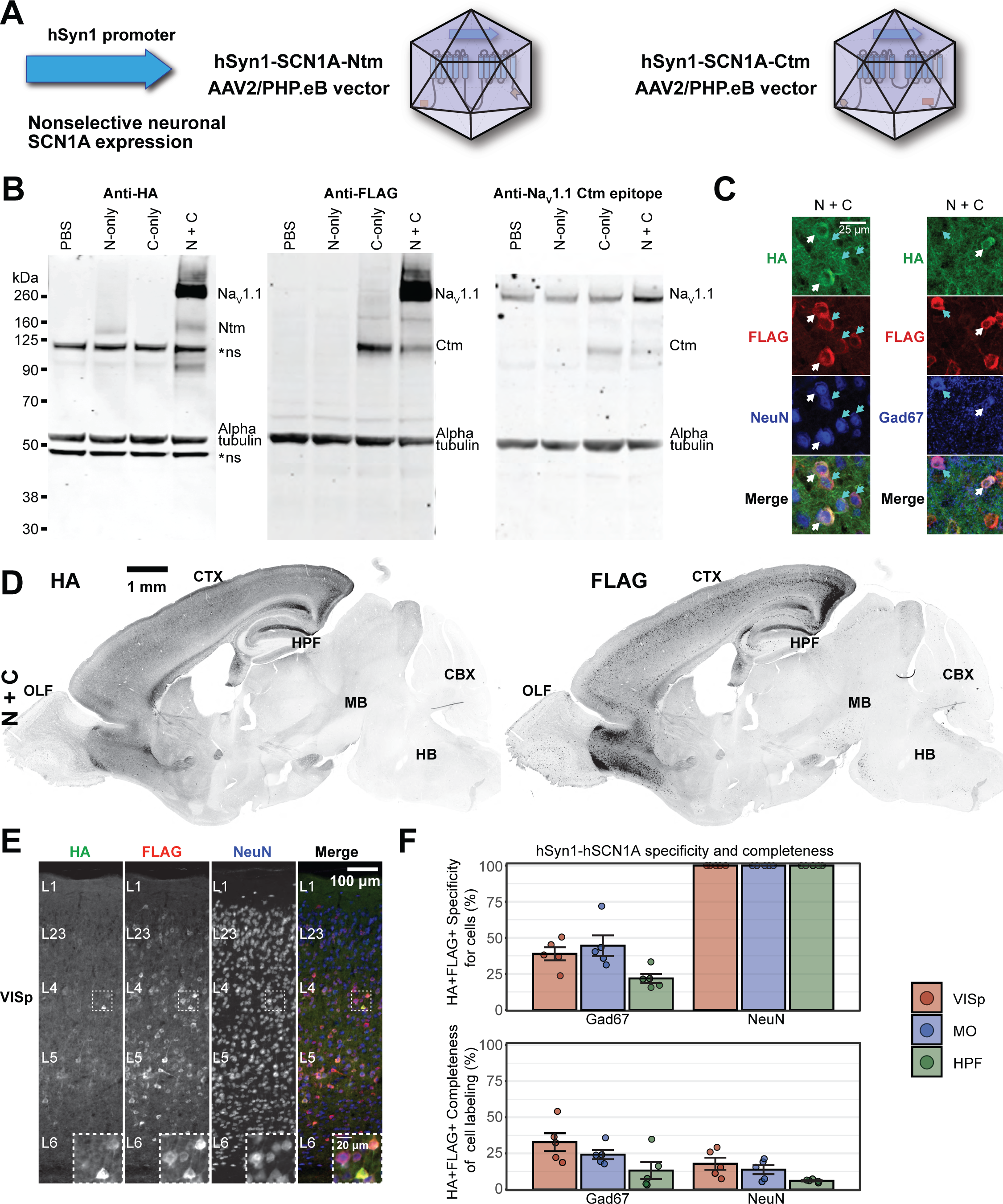
Nonselective delivery of SCN1A to neurons using hSyn1 promoter. (A) Recombinant AAV2/PHP.eB vectors for delivery of hSyn1-split-intein-SCN1A. (B) Efficient SCN1A reconstitution in mouse brain with hSyn1-split-intein-SCN1A vectors. In panels B-F we injected P2 neonatal mice BL-ICV with 3e10 gc of each indicated vector (6e10 total gc in the N+C animals). At P24 we analyzed mouse brain membrane protein fractions by western blotting with antibodies targeting HA or FLAG epitope tags, the C-terminus of Na_V_1.1, and alpha-tubulin as a loading control. Note the PBS-injected negative control lane is the same lane as that shown in Fig. 2B as these experiments were performed together. (C) HA- and FLAG-expressing NeuN+ and Gad67+ cells in cortex after co-injection with N- and C-terminal vectors. White arrows indicate NeuN+ and Gad67+ cells that express FLAG but not HA. Cyan arrows indicate NeuN+ and Gad67+ cells that express both FLAG and HA. Layer 2/3 of VISp is shown at P84. (D) Representative stitched fluorescence image of biodistribution of biodistribution of HA and FLAG epitopes in neurons after co-injection with N- and C-terminal vectors. Expression is pseudo-colored black. Expression shown at P84. (E) Representative stitched fluorescence image of HA and FLAG epitopes and NeuN+ neurons throughout the layers of VISp. Expression shown at P84. (F) Quantification of specificity and completeness of expression within Gad67+ and NeuN+ neurons in multiple telencephalic regions. We counted cells that express both HA and FLAG epitopes in VISp, MO, and HPF (layer 1 was excluded from VISp and MO analysis due to hSyn1-PHP.eB vectors inefficiently targeting that layer). Each point represents one mouse, bars represent the means, and error bars represent standard error of the mean. As expected hSyn1-driven expression shows specificity for NeuN+ cells but not Gad67+ cells. Mice span ages P76-P86, mean age P81. Abbreviations: CTX cerebral cortex, OLF olfactory areas, HPF hippocampal formation, STR striatum, MB midbrain, HB hindbrain, CBX cerebellar cortex, VISp primary visual cortex, MO motor cortex.

### Dual hSyn1 nonselective neuronal AAV vectors led to pre-weaning mortality

To characterize the efficacy of nonselective neural SCN1A AAVs to prevent SUDEP, we conducted spontaneous mortality surveillance in *Scn1a^fl/+^;Meox2-Cre* DS model mice after BL-ICV injection of dual N+C AAVs at P0-3, as compared to untreated or empty or single part control animals (**Fig. 7A**). During the preweaning weeks, mice treated with dual nonselective AAVs exhibited a surfeit of unexpected deaths. The extent of pre-weaning mortality by P21 was dose-dependent and significantly greater than that observed under negative control conditions (low dose 1e10 gc each vector, n= 9/36 [25%] deaths, Fisher’s exact test untreated comparison p= 3.6e-4; high dose 3e10 each vector, n= 8/18 [44%] deaths, Fisher’s exact test untreated comparison p= 6.6e-6, **Fig. 7B**). Since it was not possible to recover the lost pups, census of DS and control mice during pre-weaning stage were estimated based on the number of DS and control mice identified after genotyping at P21 in this experiment and the number of DS and control mice observed in our colony in untreated mice in the prior 6 months period. This analysis did not reveal any detectable influence of genotype on nonselective hSynp1 SCN1A-induced mortality during the preweaning period (**Fig. 7C**), and pathology of surviving nonselective SCN1A-expressing brains suggests no obvious microgliosis but greater DS-associated astrogliosis (**Fig. S5**). In the post-weaning period, we did not observe significant effects on survival or average age of death with either the high-dose or low-dose nonselective SCN1A AAV treatments as compared to the untreated or control AAV-treated mice (**Fig. 7B**). Together these findings indicate that nonselective neuronal expression of SCN1A offers little protection from SUDEP and concerningly, has a dose-dependent mortality side effect during the pre-weaning period.

**Figure 7:**
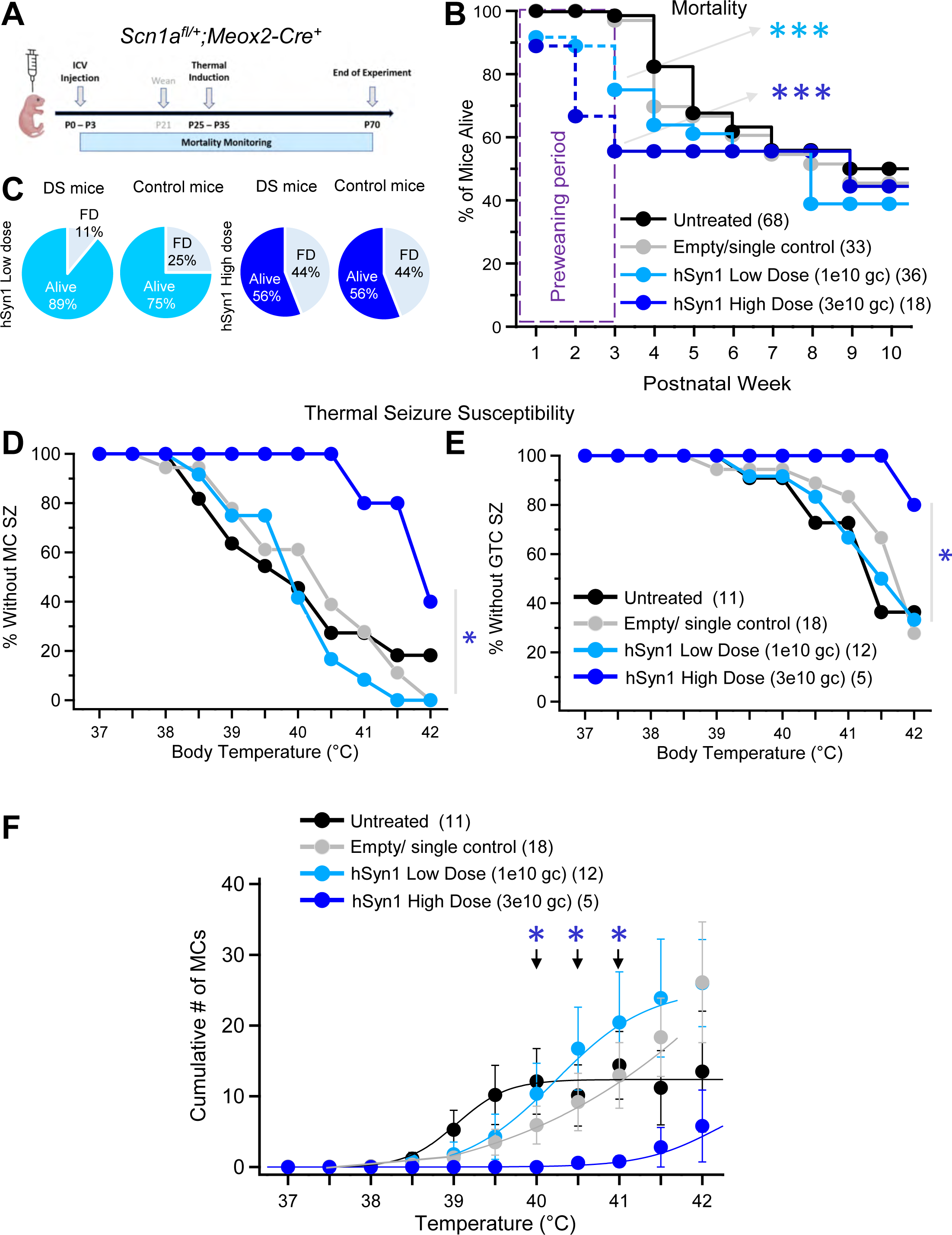
Early pre-weaning toxicity and weak protection from epileptic symptoms with nonselective SCN1A vectors. (A) Experimental timeline to rescue of epileptic symptoms in DS model mice with nonselective hSyn1 promoter-driven SCN1A vectors. (B-C) Preweaning mortality in DS model mice after treatment with nonselective SCN1A AAVs. (B) Mice treated with hSyn1 N+C SCN1A AAVs at high dose (3e10 gc each vector, n= 18) or low dose (1e10 gc each vector, n= 36) exhibit significantly greater preweaning mortality by P21 than untreated mice (n= 68) and empty/single part negative control mice (n= 33). *** p < .001 at P21 timepoint versus untreated by Fisher’s exact test. Low dose versus untreated p= 3.6e-4; low dose versus empty/single p= 0.014; high dose versus untreated p= 6.6e-6; high dose versus empty/single p= 4.9e-4. We did not observe significant effects on survival in the post-weaning period. Note the untreated control groups in **Fig. 7B, D-F** represents the same sets of untreated animals as that shown in **Fig. 3B-E**. (C) From analysis of recovered genotypes at P21 we inferred that both DS and littermate control animals were similarly affected by nonselective SCN1A AAV lethality. FD: found dead. (D-F) Protection from heat-induced seizures in DS model mice after treatment with high-dose nonselective SCN1A AAVs. (D) DS model mice treated with high-dose hSyn1 N+C SCN1A AAVs (n= 5) are significantly less likely to exhibit MC seizures by 42°C than Empty/single part mice (n= 18, * Log-rank test p= 0.020). (E) DS model mice treated with high-dose hSyn1 N+C SCN1A AAVs exhibit a trend towards less GTC seizure likelihood by 42°C than empty/single part mice (* Log-rank test p= 0.047). (F) Mice treated with high-dose hSyn1 N+C SCN1A AAVs exhibit significantly fewer MC events during thermal challenge assay than mice treated with empty or single-part vector controls (* p < 0.05, unpaired t-test at each indicated temperature for High-dose versus Untreated or Empty/single animals). In D-F, we did not observe a protective effect against seizures from low-dose hSyn1 N+C AAVs (n= 12).

### Dual hSyn1 AAV vectors confer partial protection against thermal MC and GTC seizures

To examine whether the nonselective AAV mediated gene therapy might counter thermal seizure susceptibility in DS mice, we also tested these mice with the thermal seizure induction protocol. In surviving mice treated with the high dose of dual hSyn1 AAVs, as compared to empty/single part negative controls we observed significant protection from thermally induced MC seizures (Log-rank test p= 0.020, **Fig. 7D**) and from thermal GTC seizures (Log-rank test p= 0.047, **Fig. 7E**), as well as greatly lessened cumulative MC seizure load during the thermal challenge (Untreated or Empty/single versus Treated, unpaired t-test p< 0.05 at each temperature from 40 to 41, **Fig. 7F**). In contrast we observed no protection from heat-induced MC or GTC seizures in mice treated with the lower dose of the dual AAVs compared to negative control DS mice (**Fig. 7CD-F**). Thus, despite the early mortality induced by nonselective neuronal SCN1A AAVs, they can offer protection from thermally induced MC and GTC seizures at the higher dose in surviving treated adults.

## Discussion

In this work we use an AAV viral vector system to deliver functional human Na_V_1.1 to pathological telencephalic GABAergic interneurons in DS model mice, using the optimized enhancer DLX2.0 to achieve high specificity. Importantly, we find not only that delivery to telencephalic GABAergic interneurons is sufficient for strong rescue of epileptic symptoms, but also that this specificity is required to deliver human Na_V_1.1 in a manner that can be tolerated. DLX2.0-SCN1A AAV vectors achieve long-term recovery of DS mortality (from 200 days to one year) in two independent DS mouse models at two independent testing sites. This robust mortality protection correlates with strong anti-seizure effect, suggesting the mechanism behind mortality rescue is seizure reduction due to resupplying Na_V_1.1 voltage-gated sodium channel activity.

### Which cell types are sufficient to rescue DS epilepsy using SCN1A gene replacement?

Mouse models of DS suggest that epileptic symptomology is driven by a disruption of the excitatory/inhibitory balance^12,24,25,27^, and congruently, patients display disrupted inhibition in the cortical microcircuit^26^. These observations have led to the hypothesis that DS epilepsy is primarily a disease of the interneurons^2,12,52^, and that possibly interneuron targeting might be an effective therapeutic strategy. Our results confirm this hypothesis. Within this cell class, several subclasses of telencephalic GABAergic interneurons may contribute to the disease. In particular, PVALB-expressing interneurons are thought to promote beneficial gamma rhythms^53,54^, possibly through their *Scn1a*-dependent fast-spiking behavior^25^. However, SST-expressing and other telencephalic interneuron populations may also contribute to the epileptic phenotype^25^. Importantly, the DLX2.0 enhancer used in this study targets both PVALB and SST and also other telencephalic interneuron populations, in both mouse and human tissue^31,41^, which may explain the strong anti-epileptic effect of DLX2.0-SCN1A.

### A unique SCN1A-targeting disease-modifying strategy for DS

Drug development to find pathway modulators that can overcome deficient Na_V_1.1 has been challenging with a few successes^6,55–58^. SCN1A upregulation has been a major goal to counteract the SCN1A haploinsufficiency seen in most DS patients. Recently, several groups have reported gene therapy approaches to upregulate SCN1A. Several studies have successfully boosted endogenous Na_V_1.1 with AAV CRISPR or zinc finger activators^11,49,59^, or with antisense oligonucleotides that increase the efficiency of Na_V_1.1 production^9,10^. Other studies have demonstrated success in delivering SCN1A, as a therapeutic replacement transgene, utilizing an adenovirus vector which offers high capacity but limited biodistribution^47,60^. The concept of *SCN1A* gene replacement or augmentation for DS is supported by genetic experiments demonstrating DS phenotypic reversibility with Na_V_1.1 re-expression^23^. Our AAV-mediated and telencephalic GABAergic interneuron-selective gene replacement strategy is unique in several ways. First, our vectors express a new copy of the SCN1A gene at moderate levels, and don’t upregulate the endogenous SCN1A allele, a strategy that could be ineffective or harmful for certain disease alleles. Second, the vectors target telencephalic GABAergic interneurons, the most essential cell type, while antisense oligonucleotides lack targeting ability. Since SCN1A is expressed in both excitatory and inhibitory neurons^61,62^, activation of the SCN1A in excitatory neurons could attenuate the corrective effect of SCN1A upregulation. Last, using AAV delivery, with the PHP.eB capsid and neonatal ICV injection, widespread viral transduction of telencephalic GABAergic interneurons is achieved. We are not aware of a vector that can give superior widespread transduction to the brain. None of the previous studies delivering or activating Na_V_1.1 using exogenous agents have demonstrated complete recovery of mortality as we have observed. Moreover, we have demonstrated robust rescue in two genetic models, at two research sites, and even in a severe *Dlx5/6-Cre*-driven knockout model. Thus, gene replacement, cell type specificity, and broad coverage of telencephalic interneurons provides a unique and highly effective treatment for DS.

### Potential and challenges for a gene therapy for DS patients

Currently there is no approved *SCN1A-*correcting treatment for DS. In this work, we demonstrate strong recovery of DS epileptic symptoms without toxicity in mouse models, which provides hope. However, there are several important outstanding questions that need answers before this approach could be applied to patients. First, using a dual vector system, it would be difficult to achieve similar biodistribution in a human brain as we have observed in mice after neonatal ICV injections with PHP.eB vectors. Achieving this level of transduction throughout the human forebrain may require advanced BBB-penetrant AAV capsids such as those currently in development^42,63–68^, or alternatively identifying the minimal brain regions that could be targeted with intraparenchymal delivery where local coverage can be high^69,70^. Lower levels of transduction with other virus batches resulted in less complete rescue of mortality, suggesting that functional recovery in patients would correlate with the achievable biodistribution.

Secondly, it is not known what the minimal level of expression per cell is required, and whether high of levels might become toxic over time in humans. Encouragingly, we do not observe signs of toxicity over one year in mouse brain, suggesting the intein fusion proteins delivered with cell type specificity may be safe. Finally, we do not know whether telencephalic GABAergic interneuron delivery might recover other non-epileptic DS comorbidities that cause severe quality of life issues, like cognitive deficits^71^, motor issues^72^, and sleep disturbances^73,74^. New regulatory elements^29–33^ that confer a broader distribution pattern extending to sub-telencephalic regions may be required to correct these problems. However, we have been impressed that expression of SCN1A in telencephalic GABAergic interneurons is sufficient to counteract all mortality, suggesting other associated brain structures that control cardiac and respiratory function act downstream of that circuit in DS.

### Final conclusion

We demonstrate proof-of-concept for AAV-mediated SCN1A gene replacement therapy in DS, which requires cell class specificity for safety and efficacy. These results and these vectors represent an important step towards a gene replacement gene therapy for DS patients, and possibly other conditions of pathologic insufficient sodium channel function.

## Methods

### Study design

In this study, we tested if *SCN1A* gene replacement was possible using a dual vector system to deliver both halves of the molecule and split-intein technology to fuse the expressed halves into a single scarless full length protein. We tested if SCN1A could be expressed in a circuit-selective fashion using enhancer-AAVs to deliver the transgenes to telencephalic GABAergic interneurons or all neurons. Then, we tested if these circuit-selective enhancer-AAVs were sufficient to correct major epileptic phenotypes of mouse models of DS. We selected three major phenotypes for testing that correspond to clinical phenotypes seen in DS: mortality, heat-induced seizures, and spontaneous seizures. In vitro studies were conducted with multiple biological replicates to demonstrate the full-length Na_V_1.1 functional channels were formed. Na_V_1.1 assembly in GABAergic interneurons was demonstrated in multiple replicates using viruses from two sources in vivo in mouse brains. Circuit specificity and completeness were demonstrated by immunohistochemistry, and full-length Na_V_1.1 assembly by western blot analysis. To test in vivo efficacy, litters of mice were treated between P0 and P3 with the dual viruses or single virus controls by ICV administration prior to genotyping. In vivo efficacy was further confirmed at a second site on a genetically distinct DS model. Animal sexes were noted but no differences in efficacy were observed. All experiments were run across multiple litters, and sufficient animals were injected and evaluated to draw statistically significant conclusions. Dual vectors produced from two sources were tested in vivo for efficacy. Researchers were not blinded to the animal genotypes after they were determined, or to vectors delivered. No data were excluded from the study. For statistical analysis, data are reported as means ± standard error of the means, Kaplan-Meier survivor plots, or grouped bar graphs. Comparisons across continuously distributed grouped data were prefaced by Shapiro-Wilk tests for distribution normality. When normality assumptions held, the groups were compared by unpaired t-tests or ANOVA, but when normality is not held, the groups were compared by Mann-Whitney U test. To compare Kaplan-Meier survivorship, we used Mantel-Cox or Log-rank tests. To compare groups of categorical count data, we used Fisher’s exact tests. Differences were considered significant at p< 0.05, with Bonferroni adjustments to significance thresholds with comparisons across multiple groups.

### Mice at Seattle Children’s Research Institute (SCRI)

For studies conducted at SCRI, all experimental procedures were conducted in compliance with the *Guide for the Care and Use of Laboratory Animals* of the National Institutes of Health and were approved by the Institutional Animal Care and Use Committee (IACUC) of the Seattle Children’s Research Institute under protocol ACU000108 (PI-Kalume). WT and mutant mice within litters were used in these experiments. They were subjected to a treatment or control paradigm as described below. Both male and female mice were included in the studies. Each litter was randomly assigned to the treatment or control group.

Mice at SCRI were maintained in standard cages for laboratory mice, on a 12h:12h light-dark cycle, with *ad-libitum* access to food and water, at 23 degrees C. Mouse models and their littermates of DS used in these studies were generated using Cre-Lox technology. DS mice carrying a whole body heterozygous knock-out of *Scn1a* were obtained by breeding floxed *Scn1a* mice^24,45^ with *Meox2*-Cre mice (Strain #: 003755; Jackson Laboratories). DS mice carrying an Scn1a KO allele restricted to (specifically in) forebrain GABAergic neurons alone were generated by breeding floxed *Scn1a* mice with *Dlx5/6-Cre* mice (Strain#: 008199, Jackson Laboratories). All breeder mice were maintained on a C57BL/6J background for at least 10 generations. Animals were genotyped for *Scn1a* floxed allele using the following primers: FHY311 (5’-CTTGATGTGTTGAAATTCAC-3’) and FHY314 (5’-TATAGAGTGTTTAATCTCAAC-3’) which yielded a 846 BP WT allele, 1019 BP floxed allele; and 258 BP excised allele. For the Cre alleles, the following primers were used: (5’-GGTTTCCCGCAGAACCTGAA-3’) and (5’-CCATCGCTCGACCAGTTTAGT-3’) (Jackson Laboratories)

### Thermal seizure test

Protocols described in our previous work were used at SCRI^46,50^. Mouse core body temperature was monitored and controlled using a rectal temperature probe (RET4) and a heat lamp, both connected to a temperature controller in a feedback loop (Physitemp Instruments Inc.). Baseline body temperature was measured and subsequently, gradual temperature increases of 0.5°C every 2 minutes were conducted until seizure occurrence or a 42°C temperature was attained. Then, the mouse was immediately cooled down using a small fan. Mouse behavior during the whole test was recorded using a digital video camera and reviewed for MC and GTC seizure scoring.

### Intracerebroventricular Injections at SCRI

A slightly modified injection technique from Kim et al. was adopted^75,76^. Neonatal P0-3 mice were cryo-anesthetized on a small aluminum plate placed on ice. Single AAVs or dual AAVs were injected bilaterally into lateral ventricles using a 33-gauge needle attached to Hamilton microliter syringe. 2.5 μl of the AAV solution were injected in each ventricle for a total of 5 μl per mouse containing a total of 1e10 or 3e10 gc each viral vector. Following the injection, mice were put back into their nest and placed on a warming pad until their body temperature returned to normal. Subsequently, they were returned to the cage with the mother.

### Electrocorticography (ECoG) electrode implantation surgery at SCRI

Mice underwent survival surgery to implant ECoG and EMG electrodes, under isoflurane anesthesia, and using similar procedures as reported in our prior work^77,78^. A midline incision was made above the skull to expose the site of electrode implantation. ECoG electrodes consisted of a micro-screw attached to a silver wire (diameter: 130 μm bare; 180 μm coated). EMG electrodes were made of a silver wire shaped in a loop at one end. An ECoG electrode micro-screw was inserted into a small cranial burr hole above the somatosensory cortex in each hemisphere. Similarly, a reference electrode micro-screw was placed in a burr hole above the cerebellum. EMG electrodes were inserted and secured into the neck muscles. All electrodes were attached to an interface connector and the assembly was affixed to the skull with dental cement (Lang Dental Manufacturing Co., Inc., Wheeling, IL, United States). The incision around the electrode implant was closed using sutures. Mice were allowed to recover from surgery for 1–3 days before recording.

### Video-ECoG-EMG recording at SCRI

Simultaneous video-ECoG-EMG records were collected in conscious mice on a PowerLab 8/35 data acquisition unit using LabChart 8.0 software (AD Instruments, Colorado Spring, Co). All bioelectrical signals were acquired at 1-KHz sampling rate. The ECoG signals were processed with a 1–70 Hz bandpass filter and the EMG signals with a 10-Hz highpass filter. Power-spectral densities of the electrical signals were computed, and video-ECoG-EMG records were inspected for interictal spikes and ictal epileptiform events. Interictal spikes were characterized on ECoG as discharges with an abrupt onset, a sharp contour, and an amplitude greater than twice the background activity. Interictal spikes were frequently followed by a slow wave, but they were not associated with increased EMG activity or movement on video. Conversely, GTC seizure events were marked at their onset by bursts of generalized spikes and waves of increasing amplitude and decreasing frequency on ECoG. They coincided with increased activity on EMG and video and were followed by a distinct period of post-ictal ECoG suppression.

### Mice at Allen Institute for Brain Science (AIBS)

For studies conducted at AIBS, all mice were handled under appropriate institutional protocols and guidelines. Procedures were approved by the Allen Institute Institutional Animal Care and Use Committee under protocols 2002 and 2301. We housed animals in a 14:10 light:dark cycle in ventilated racks with ad libitum access to food (LabDiet 5001) and water, as well as enrichment items consisting of plastic shelters and nesting materials. Young animals are weaned promptly at 21 days of age. We obtained *129S1/SvImJ-Scn1a^em1Dsf^/J* (here *Scn1a^+/R613X^*) mice from Jackson Laboratory (strain # 034129) and maintained breeders on a 129S1/SvImJ genetic background. To generate experimental animals to model DS, we crossed these animals with C57Bl/6J mice (Jackson strain # 000664), which resulted in a 50:50 129:BL/6 F1 genetic background which has been successfully used to model epileptic phenotypes of DS in animals containing one loss-of-function allele of *Scn1a* ^48,60,79,80^. Genotyping was performed with tail biopsy at P2, and we utilized PCR-sanger genotyping services at Transnetyx for this line. For survival monitoring we checked cages twice daily for the presence of deceased animals.

### Neonatal ICV injection at AIBS

We used the neonatal ICV injection technique of Kim et al.^76^. Briefly, we anesthetized P2 neonates with ice but shielded from direct ice exposure. During anesthesia, pups were injected freehand bilaterally with 5 μL (2.5 μL each hemisphere) of AAV-containing solution using a Hamilton syringe. AAVs were diluted in sterile PBS to expel either 1e10 or 3e10 gc of each of two halves of the dual-vector encoding split-intein human SCN1A. In control animals, only one half of the dual-vector system was delivered, or DLX2.0-SYFP2-only empty control vector (CN1390^31^), or mice were left untreated. Control vectors were delivered at 3e10 gc per animal with BL-ICV delivery. After injection, pups were gently warmed on a cage warmer set to 28°C with mother present.

### Continuous video ECoG/EMG recordings at AIBS

We implanted adult mice (P56-90) with ECoG/EMG headmount similarly as previously described^81^. For stereotaxic surgical procedures, we induced anesthesia in mice first with 5% isoflurane in oxygen, and then maintained anesthesia with 1.5-2.5% isoflurane. We implanted screw electrodes with wire lead (0.08”, #8405, Pinnacle Technology Inc., KS, USA) over the left somatosensory cortex (AP: + 1 mm; ML: - 2.5 mm), the right parietal lobe (AP: - 2 mm; ML: + 1.5 mm), the right frontal area (AP: + 1 mm; ML: + 2.5 mm; as Ground), and the cerebellum (AP: - 5.8 to -6.2 mm; ML: 0 mm; as Reference). Electrode leads were soldered onto the 8-pin headmount (#8431-SM, Pinnacle Technology Inc.). The headmount contains two insulated EMG wire electrodes that are pre-soldered, and these EMG electrodes were inserted into the neck muscles. All wires, pins and the headmount were embedded in light curable dental composite resin (Prime-Dent, Prime Dental Manufacturing Inc., Chicago, IL, USA). Mice were singly housed post-surgery and recovered for at least 7 days prior to recording. Recordings were thus acquired between ages P74 and P123.

For recordings at AIBS, mice were singly housed in 10-inch clear acrylic chambers (#8228, Pinnacle Technology Inc.) under a 14-hr on, 10-hr off light/dark cycle. Mice were tethered with the pre-amplifier through a commutator to the data acquisition system (#8401-HR, Pinnacle Technology Inc.). All ECoG/EMG data were recorded with a 500 Hz sampling rate, 10 X gain, a low pass (ECoG: 0.5 Hz; EMG: 1 Hz) filter, and a high pass (500 Hz) filter. Videos were recorded synchronously at a frame rate of 10 frame/s with a resolution of 640x480 pixels. We implanted a total of 18 non-injected *Scn1a^+/R613X^*mice. Of these 18, seven mice (3M+4F) died during recovery prior to recording, and we recorded from the remaining 11 mice (9M+2F). For these non-injected animals we recorded for 53 to 335 hrs, with some recordings prematurely shortened because of death during the recording session. We also implanted a total of 11 DLX2.0-intein-SCN1A-injected *Scn1a^+/R613X^*mice. Of these 11, we could not record from three mice due to surgical error (1M), hardware failure (1M), and one death following surgery (1F), and we recorded from the remaining 8 mice (4M+4F). For these DLX2.0-intein-SCN1A-injected *Scn1a^+/R613X^* mice we recorded animals for 207-257 hrs.

To quantify ECoG data at AIBS, we quantified the number of GTC seizures with manual counting over the recording period and expressed as the average number of GTC seizures per 24 hrs of recording. We also counted the number of interictal spikes (IISs) during the last 24 hrs of recording for each mouse. To do so, we identified candidate interictal spike events by cumulative line length over a time interval of 50 msec with a threshold of 50 microvolts, as previously described^74,82^, plotted each candidate event for visual confirmation of spike-like characteristics (high amplitude strong deflection and return to baseline within 30 msec) and counting. These measurements of epileptic activity are non-normally distributed among animals by Shapiro-Wilk tests for normality (GTCs within non-injected *Scn1a^+/R613X^* mice: W= 0.81, p= 0.018; IISs within non-injected *Scn1a^+/R613X^* mice W= 0.72, p= 0.0015), we compared number of seizures and IISs were compared between groups by two-sample Mann-Whitney U test (with significance level at p= 0.05).

### IHC

Under avertin terminal anesthesia, we perfused mice with ice-cold PBS with 0.25 mM EDTA added (25 mL), followed by cold 4% PFA in PBS (12 mL). Following brain and other organ dissection, we post-fixed brain in 4% PFA in PBS overnight at 4°C. We prepared PFA in PBS in one liter-sized batches by dissolving PFA powder in PBS with heating, and froze 50 mL aliquots at -20°C until use, which was important as we found anti-Gad67 and anti-GABA stain quality depended upon PFA preparation method. After overnight postfixation, we transferred brains to 30% sucrose solution in PBS, and then embedded in OCT after sinking (48-72 hours), froze on dry ice and stored at -80°C until sectioning. We sectioned brains sagittally at 25 micron thickness using a Leica CM3050S cryostat and stored sections in PBS at 4°C until IHC. For IHC we permeabilized and blocked sections with blocking solution (PBS containing 5% normal goat serum [Thermo Fisher Scientific # 10000C] and 0.1% Triton X-100 [Millipore-Sigma # X100-100ML]) for 60 minutes. Then we probed with diluted primary antibody in blocking solution overnight, washed twice for 15 minutes each with PBS containing 0.1% Triton X-100 (PBSX), then detected with diluted 488-, 555-, and 647-conjugated secondary antibodies (all from Thermo Fisher Scientific) along with DAPI (4′,6-Diamidino-2-phenylindole dihydrochloride, Millipore-Sigma # 10236276001, used at 1 μg/mL), washed twice with PBSX for 15 minutes each, then mounted sections on SuperFrost Plus slides (VWR # 48311-703) in Prolong Gold (Thermo Fisher Scientific # P36930), and imaged after overnight curing. We imaged slides on a Nikon TI-Eclipse epifluorescent or an Olympus FV-3000 confocal microscope.

We used the following primary antibodies: mouse monoclonal anti-FLAG clone M2 (Millipore-Sigma # F1804), rabbit monoclonal anti-HA clone C29F4 (1/1000, Cell Signaling # 3724S), mouse monoclonal anti-HA clone 16B12 (1/1000, Biolegend # 901513), mouse monoclonal anti-HA clone HA.C5 (1/1000, Thermo Fisher Scientific # MA5-27543), mouse monoclonal anti-Gad67 clone 1G10.2 (1/250, Millipore-Sigma # MAB5406), mouse monoclonal anti-NeuN clone 1B7 (1/500, Novus Biologicals # NBP1-92693AF647), guinea pig polyclonal anti-GABA (1/500, Millipore-Sigma # AB175).

For costains with anti-Gad67, we omitted Triton X-100 detergent permeabilization during Gad67 immunoprobing and detection, then re-fixed antibodies onto sections with 4% PFA in PBS for 15 minutes at room temperature, then washed with PBS twice for 15 minutes, then performed a second round of permeabilization and reblocking and antibody staining with desired co-staning antibodies, as previously described^83^. This technique permits the best detection of Gad67+ inhibitory neurons.

### SCN1A ORF design, cloning, and packaging

To design full-length human SCN1A for delivery in split intein-fusion SCN1A halves we started with the 1998-amino acid RefSeq sequence NP_001340878.1. This is because previous single cell RNA-seq studies in mouse and human^61,62^ demonstrate the major species expressed by cortical cell types includes the 348-bp form of the 11^th^ exon (of 26 exon numbering scheme), which corresponds to a full open reading frame of 1998 amino acids. The 2009 amino acid isoform is expressed at a lower level across multiple cell types including PVALB neurons (see Supplementary Figure 1A-B). We also included a threonine at position 1056 of 1998 (corresponding to position 1067 of the 2009-amino acid isoform). This residue is alanine in the NCBI RefSeq sequence but is a threonine in commercial clones available from Origene (catalog # RG220167), as well as a conserved threonine across most other mammalian species (**Fig. S1C**), and finally we observed this residue to be a threonine in three of three human tissue donors sequenced in our previous work^31^ (**Fig. S1D**, data available at dbGaP # phs002292.v1.p1). From gnomAD^84^ the reference alanine allele appears to be the minor allele in the human population (27%), whereas the majority of alleles in the population encode threonine at that position (73%). As a result, we used threonine in that position for our SCN1A transgene.

From this protein sequence we split the protein into two halves at the natural cysteine at position 1050 (numbering according to 1998-amino acid isoform), and we appended the Cfa-N and Cfa-C intein proteins to the ends of the split breakpoints with no linkers, so that the natural cysteine residue would work as the C+1 extein and result in scarless protein joining. The native Npu split-intein prefers hydrophobic residues prefers hydrophobic residues such as the native methionine at the C+2 position^36,85^, which we reasoned would likely promote half joining efficiency. We also appended HA and FLAG epitope tags (HA at the N-terminus of the N-terminal half, and FLAG at the C-terminus of the C-terminal half), with short dipeptide linkers between the epitope tags and SCN1A coding sequence. With these protein sequences we reverse-translated and performed codon optimization using Integrated DNA Technologies online codon optimization tool (https://www.idtdna.com/pages/tools/codon-optimization-tool), and manually adjusted the codon usage to minimize cryptic splice donors and acceptors which could negatively impact expression from unwanted splicing, and manually minimized repeats over 10 bp which might negatively impact cloning or expression. The full protein open reading frame (ORF) was 100% identical at the amino acid level to the native isoform, but only 76.6% identical at the nucleotide level. We also inserted a human intron from hemoglobin which we engineered to reduce possible cryptic TATA-box promoter sequences, as well as produce efficient splice donor and acceptor sites without need for exonic splice enhancer sequences. We then synthesized these sequences as G-blocks (Integrated DNA Technologies) and cloned them using Infusion kit (Takara biosciences catalog # 638948) into pSMART-HC-Kan (Lucigen) vectors along with upstream CMV promoters and downstream IRES2-SYFP2 or IRES2-mScarlet transfection reporters for expression in HEK-293 cells. Alternatively, for full-length human SCN1A we assembled synthetic Gblocks lacking the intein fusion proteins but with overlapping ends using Infusion kit into pSMART alongside CMV promoter and downstream IRES2-mScarlet reporter. The full length human SCN1A pSMART vector contained only a C-terminal FLAG epitope but not an N-terminal HA epitope. For cloning into pAAV vectors we inserted the intein-fusion halves into CN1390 (Addgene plasmid #163505) in place of SYFP2 reporter for DLX2.0-driven expression or into CN1839 (Addgene plasmid #163509) in place of SYFP2-10aa-H2B for hSyn1-driven expression. During this cloning we also replaced BGH polyA in the original vectors sequences with shorter synthetic polyA sequences due to size constraints^86^.

Propagating full-length SCN1A sequence-containing plasmids in bacteria is challenging^16,18,39,87^. To minimize rearrangement of full length SCN1A we used pSMART-HC-Kan backbone (Lucigen) which helps minimize backbone-driven transcription in bacteria. We also propagated SCN1A into CopyCutter EPI400 cells (formerly Lucigen, now BioSearch Technologies, catalog # C400CH10) for low-copy growth in absence of CopyCutter Induction Solution, requiring larger culture volumes to compensate for reduced plasmid yields (5 mL for minipreps, 0.5-1L for maxipreps). Due to their low-copy nature, the full length SCN1A plasmid preps from these CopyCutter EPI400 cells show high contamination from bacterial genomic DNA. We minimized this contamination by adsorbing flocculated genomic DNA against copper-chelated agarose resin^88^ (G Biosciences catalog # 786-285), followed by repurification and concentration on Ampure XP beads (Beckman-Coulter # A63882). Each batch of full-length SCN1A plasmid required full sequence validation by Sanger sequencing across the complete ORF to verify absence of rearrangement prior to use, in the case of minipreps using PCR amplification of the full ORF to generate enough DNA for sequencing. Alternatively, large-scale maxipreps of full-length SCN1A for transfection experiments were produced by Aldevron (Fargo, ND) using their proprietary growth conditions, which were confirmed to be fully intact by full ORF sequencing. In contrast to the special challenges experience with full length SCN1A ORF, plasmids containing the split fusion protein halves alone did not exhibit rearrangements or require special culturing techniques, and we amplified these plasmids in either pSMART-HC-Kan or pAAV backbone using Stbl3 cells (Thermo Fisher Scientific # C737303). For in house preps, all bacteria were grown at 32°C with either 50 μg/mL ampicillin (liquid growth, Millipore-Sigma # A8351-5G), 50 μg/mL carbenicillin (agar plates, Teknova # L1008), or 25 μg/mL kanamycin (liquid growth, Millipore-Sigma # K1377-5G, or agar plates, Teknova # L1023). Liquid growth was performed in a 50:50 mix of LB and TB (Teknova L8000 and T7060). Additionally, we also cloned a recombinant codon-optimized hSCN1B-P2A-hSCN2B ORF to promote folding and activity of Na_V_1.1 channels^16,17^. Packaging into PHP.eB particles with iodixanol gradient purification was performed in-house as previously described^31^, or else performed commercially at PackGene, Inc. (Guangzhou, China, and Houston, TX, USA).

### Western blot

To prepare protein samples from HEK-293 cells, we plated HEK-293 cells (ATCC catalog # CRL-1573) with fewer than 15 passages onto 12-well plates in HEK-293 growth medium (DMEM [Gibco # 10566-061] with 10% FBS [Gibco #16140-071] and 1x Pen Strep [Gibco #15070-63]) and transfected them when 50-70% confluent. We transfected 1000 ng of DNA per well using PEI-MAX (Polysciences # 24765-100) using a ratio of 1000 ng of DNA : 5μl 1 mg/mL PEI-MAX in Opti-MEM (Gibco #11058-021). For expression of SCN1A or split SCN1A halves, we also co-transfected a CMV-hSCN1B-P2A-hSCN2B construct at a ratio of 900 ng alpha subunits: 100 ng beta subunits. We replenished the medium at 12-18 hours post-transfection with fresh HEK-293 growth medium, and then harvested the cells at 48-72 hours post-transfection. For harvesting we rinsed with PBS and then lysed with RIPA buffer (Pierce #89901) containing 1x Halt Protease Inhibitor Cocktail (Thermo Fisher # 87786).

To prepare mouse brain membrane protein samples, we dissected motor cortex samples (∼25-50 mg wet tissue) into 1.7 mL Eppendorf Lo-Bind tubes and froze on dry ice and stored at - 20°C. We prepared membrane protein samples using the Mem-Per Plus Membrane Extraction Kit (Thermo Fisher #89842) with small adjustments to the protocol. Briefly we washed the tissue once with ice-cold Cell Wash Solution, spun down, and wash buffer aspirated off. Then we permeabilized tissue with 200 μL ice-cold Permeabilization Buffer supplemented with 1X Halt Protease Inhibitor Cocktail, pipetted fifty times with a Rainin P200 pipet, tumbled at 4°C for 15 minutes, and centrifuged at 18,000g for 15 minutes at 4°C. This supernatant was saved as cytoplasmic protein fraction, and we solubilized the pellet containing membrane proteins in 200 μL Solubilization Buffer with pipetting fifty times again with the Rainin P200 pipet and tumbled at 4°C for 15 minutes. Finally, we centrifuged the samples again (18,000g for 15 minutes at 4°C) and saved the supernatant as membrane protein fraction. Aliquots of this membrane protein were diluted with five volumes of PBS prior to protein quantification using BCA assay.

For western blots of HEK-293 cells and mouse brain membrane proteins, we quantified protein samples using BCA assay kit (Thermo Fisher Scientific # 23225) and analyzed them on a plate reader (Perkin Elmer/EnSpire 2300). For each lane we treated 15-20 μg protein samples with 4X NuPAGE LDS Sample Loading buffer (Thermo #NP0007) and 5% 2-mercaptoethanol (Sigma #M3148-25ml), heated them at 70°C for 6 minutes, chilled them on ice, then separated them on a NuPAGE 4-12% Bis-Tris gel (Thermo Fisher Scientific # NP0323BOX) using NuPage MOPS SDS running buffer (Thermo Fisher #NP0001) at 90V for 2 hours, alongside 5 μL Chameleon Duo Pre-stained Protein Ladder (LiCor # 928-60000) as a sizing standard lane. We then transferred the proteins to nitrocellulose membranes with pre-chilled NuPage Transfer Buffer (Life Technologies #NP0006-1) at 90V for 2 hours on ice.

After transfer, we blocked membranes with 5% milk in PBS for 1 hour at room temperature and probed overnight with rocking at 4°C with the following primary antibodies: mouse anti-FLAG clone M2 (1/1000 dilution, Sigma # F1804), mouse anti-HA clone 16B12 (1/3000 dilution for HEK-293 cell lysates, Biolegend # 901513), rabbit anti-HA clone C29F4 (1/1000 dilution for brain membrane protein preparations, Cell Signaling # 3724S), NeuroMab mouse anti-Na_V_1.1 clone K74/71 (1/300 dilution, Antibodies Incorporated # 75-023), and loading control rabbit polyclonal anti-alpha tubulin (1/5000 dilution, Cell Signaling #2144S) or mouse anti-alpha tubulin clone DM1A (1/5000 dilution, Santa Cruz Biotechnology sc-32293) in 2.5% milk in PBS with 0.1% Tween 20 (PBST). Following primary antibody, we washed 3x with PBST for 10 minutes each, then performed secondary antibody detection using appropriate IRDye 680LT- or 800CW-labeled goat secondary antibodies (LI-COR) in PBST for 1 hour, then washed 3x with PBS for 10 minutes each, then imaged the blots on the LI-COR Odyssey imager. We performed all antibody and wash steps at room temperature with gentle agitation.

### Electrophysiology

HEK293 cells (CRL-1573; ATCC, Gaithersburg, MD) were cultured in standard media consisting of DMEM, high glucose, glutaMAX (Gibco 10566016; Thermo Fisher Scientific, Waltham, MA), supplemented with 10% (v/v) fetal calf serum (FCS) (Gibco A5670401; Thermo Fisher Scientific, Waltham, MA) and 1% (v/v) Penicillin-Streptomycin (P/S) (10,000 U/ml) (Gibco 15140148; Thermo Fisher Scientific, Waltham, MA), grown at 37°C and 5% CO_2_. Cells were passaged in T25 tissue culture flasks (FB012935; Thermo Fisher Scientific, Waltham, MA) approximately twice a week. Only cells passaged less than 20 times were used for transfection and expression studies.

Plasmid constructs were acutely transfected into HEK293 cells using Viafect reagent (E4981; Promega, Madison, WI), following the manufacture’s protocol. In brief, HEK293 cells were first prepared for transfection by plating into 12-well tissue culture plates (Nunc 12-565-321; Thermo Fisher Scientific, Waltham, MA) at a density of ∼0.5-2 x10^5^ cells per well and grown to ∼80-90% confluence with standard media, allowing for one confluent well per transfection condition. On the day of transfection, media in confluent wells to be transfected were replaced with 0.5 mL fresh DMEM with 10% FCS, without P/S. Lipophilic/DNA transfection complexes were generated for each well to be transfected. This was achieved by combining a total of ∼1.0 μg of plasmid DNAs with serum-free OptiMEM (Gibco 31985062; Thermo Fisher Scientific, Waltham, MA) to a final volume of 100 μL, then adding 3.0 μL Viafect with gentle trituration and allowing the mixture to assemble at 24°C for 30 mins. After 30 mins, this mixture was added to each well. Live transfected cells were incubated overnight at 37°C, and visually monitored for transfection efficiency *in situ* using a plate microscope equipped with fluorescence (Invitrogen EVOS M7000; Thermo Fisher Scientific, Waltham, MA). Transfection efficiencies were typically >70-80%.

Specific amounts of plasmid DNAs (pDNAs) used for transfections per well: a) Full-length SCN1A (SCN1A-FL) 0.8 μg of pDNA, b) SCN1A-Ntm (SCN1A-N) 0.4 μg pDNA combined with SCN1A-Ctm (SCN1A-C) 0.4 μg pDNA, c) SCN1A-Ntm (SCN1A-N) 0.8 μg pDNA, d) SCN1A-Ctm (SCN1A-C) 0.8 μg pDNA, e) empty vector (SYFP only) 0.8 μg pDNA. All transfection conditions (a-e) were performed in a background of 80 ng pDNA of a bi-cistronic construct expressing SCN1B and SCN2B (β1/2), which encodes two sodium channel β-subunits, yielding an α:β subunit-encoding pDNA mass ratio of 10:1. One additional control utilized only 80 ng of β1/2 pDNA.

Following overnight incubation, cells in transfected wells were dissociated with TrypLE (Gibco 12-604-013, Thermo Fisher Scientific, Waltham, MA); and replated at low density onto 12 mm poly-D-lysine-coated glass coverslips (GG-12-pdl; NeuVitro, Vancouver, WA) in 24-well tissue culture plates (FisherBrand FB012929; Thermo Fisher Scientific, Waltham, MA), for patch-clamp electrophysiology. Typically, ∼10,000-15,000 cells were replated per well at sufficiently low density to isolated individual cells. This was necessary to prevent the formation of electrical junctions between contacting cells, which precludes adequate space-clamp recording conditions. Recordings were performed from 0.5-3 days after replating at low densities.

For patch-clamp recordings, coverslips containing adherent transfected cells were transferred to the stage of a Zeiss AxoExaminer.A1 microscope, equipped with an 40X water immersion objective and epifluorescence capability. Pipettes were positioned with a Sutter MPC-325 micromanipulator (Novato, CA). Whole-cell voltage-clamp recordings were acquired with an AxoClamp200B amplifier (Molecular Devices, Union City, CA), using pClamp10.4. The composition of recording solutions followed Jiang, *et al.*^89^: Bath (in mM, 140 NaCl, 2 CaCl2, 2 MgCl2, 10 HEPES, pH 7.4); Pipette internal solution (in mM, 35 NaCl, 105 CsF, 10 EGTA, 10 HEPES, pH 7.4). Patch pipettes were pulled from borosilicate glass (1B120F-4; World Precision Instruments, Sarasota, FL) on a P-97 Sutter Instruments puller (Novato, CA), and fire-polished on a Micro-Forge MF-830 (Narashige International USA, Amityville, NY) to a resistance of 0.8-1.5 MΩ. Currents were allowed 5-10 mins to stabilize after achieving whole-cell recording configuration, and acquired at 50 kHz, filtered at 5 kHz. Capacitive transients were subtracted using a P/4 subtraction scheme^90^, employing a current subtraction template derived from scaling the voltage command protocol by one-fourth. Series resistance compensation was >90% for all recordings. For voltage-dependent measurements of activation (G/V) and steady-state inactivation (SSI) which require optimal voltage-clamp conditions, currents larger than 6 nA were excluded. All currents were included for peak current density measurements. Additional initial recordings were obtained with an Axopatch 1D amplifier, which provided data only for peak current density analysis.

For conductance/voltage (G/V) plots, conductance (G) was calculated by the formula:

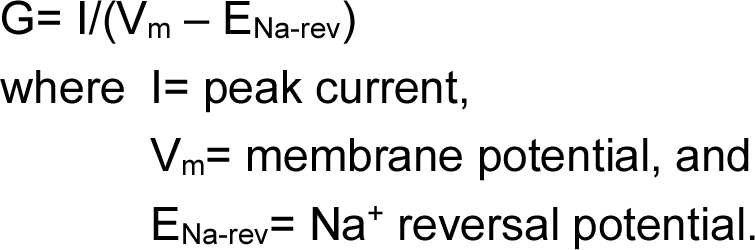

Na^+^ reversal potential was 35 mV, based on a calculated Nernst equilibrium potential with the recording solutions used.

Peak currents were recorded in response to a family of voltage steps from a holding potential of-120 mV to 40 mV, in 5 mV increments, with an inter-pulse interval of 2 seconds to allow channels to fully deactivate to the deep closed state. Conductance/voltage plots were fitted to a single Boltzmann function:

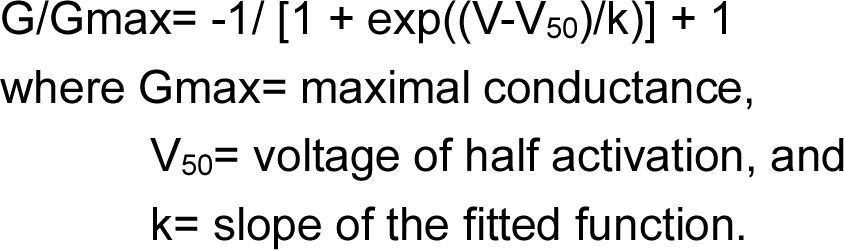

For steady-state inactivation (SSI) plots, residual peak currents activated by a step to -20 mV were measured, after a family of 1 second activating preconditioning voltage steps from -120 mV to 30 mV, in 5 mV increments. Peak residual currents after steady-state inactivation were plotted and similarly fitted to a single Boltzmann function.

Current traces were analyzed and plotted using pClamp10.4 (Molecular Devices, San Jose, CA) and Origin 8.5 (Northampton, MA). All G/V and SSI data were plotted as means with standard errors (SE) using Origin 8.5. Current density plot and all statistical calculations were performed in Prism (GraphPad, La Jolla, CA). Figures composed in Microsoft PowerPoint (Redmond, WA).

## Ethics Declarations

Competing interests

JKM, FKK, BPL, JTT, ESL are named inventors on patents related to this work. BPL is a scientific advisor for Patch Bioscience.

## Acknowledgements

We would like to acknowledge Kathryn Gudsnuk and Bargavi Thyagarajan for programmatic support. This work was supported by the following grants: RF1MH114126-01 from the National Institute of Mental Health to BPL, JTT, and ESL; UG3MH120095-01, -02, -03 from the National Institute of Mental Health to BPL, JTT, ESL, and FKK, and UF1MH128339-01 from the National Institute of Mental Health to BPL and JTT. We also would like to acknowledge the estate of Paul G. Allen for his vision, encouragement, and support.

---

**Supplementary Figure 1:**
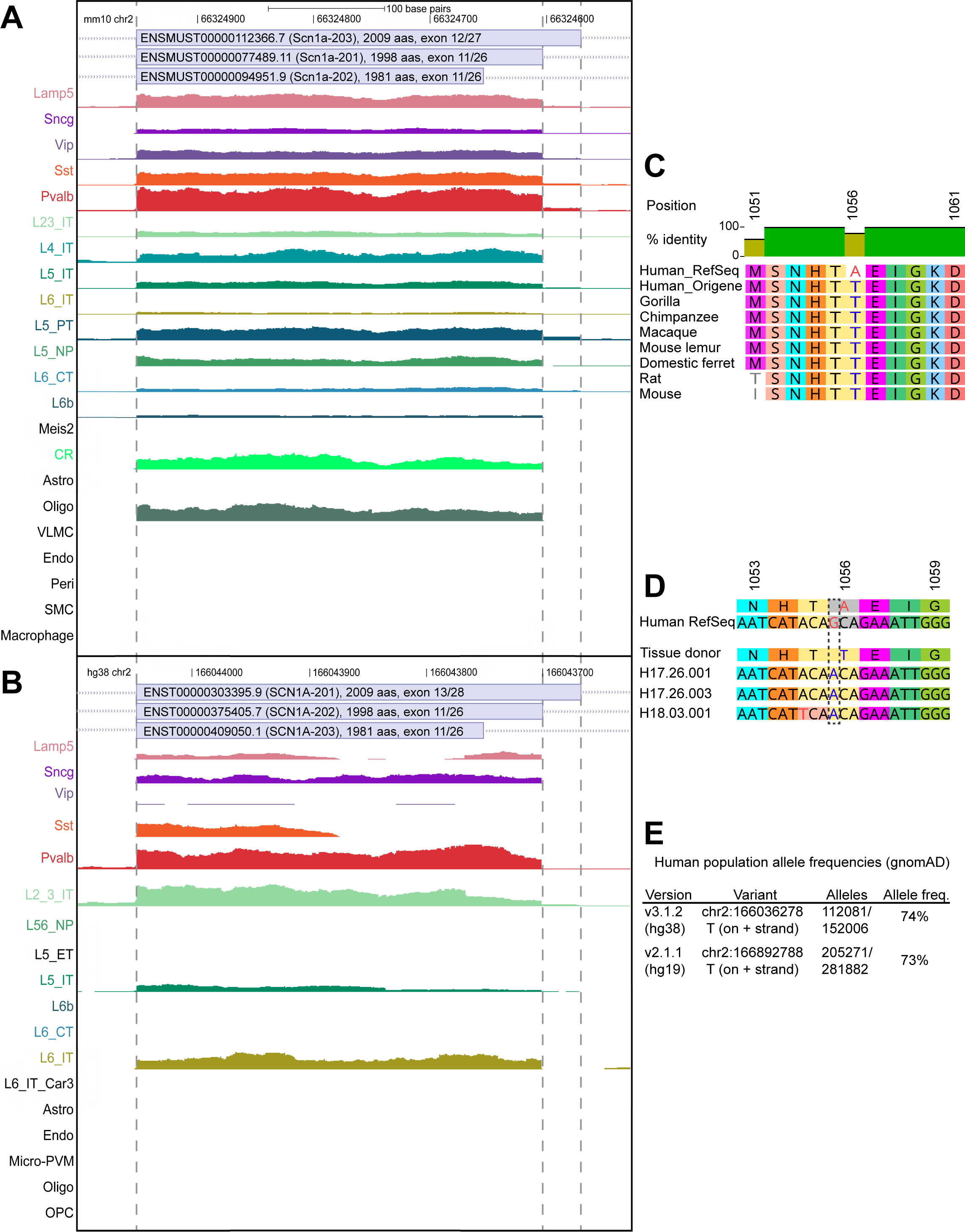
Isoform usage and allele prevalence of human *SCN1A*. (A-B) SCN1A isoform usage across cortical cell type subclasses in mice (A) and humans (B). Mouse VISp cortical cell type-specific RNA-seq profiles are from Tasic, et al.^62^ and human middle temporal gyrus (MTG) cortical cell type-specific profiles are from Hodge, et al.^61^ Genome-aligned reads are aggregated according cell type subclasses, and visualized as pileups on UCSC genome browser alongside the positions of the exon whose splice donor usage determines whether the 2009-, 1998-, or 1981-amino acid isoform of SCN1A is expressed. Regions shown: mm10 chr2:66324527-66325003, hg38 chr2:166043623-166044099 (reverse complement reference sequences for legibility). Full vertical scale represents 0.65 (mouse) or 0.4 (human) read counts per million. (B) Alignment of mammalian SCN1A protein sequences. The alanine residue at position 1056 in the NCBI RefSeq human SCN1A sequence is orthologous to a conserved threonine residue in most other mammalian species. Additionally, Origene commercial clones of human SCN1A contain threonine residue at this position, but agree at all other positions. Sequences used for alignment: human NP_001340878.1, gorilla XP_055236229.1, chimpanzee XP_054535897.1, Macaque XP_001101023.1, mouse lemur XP_020143996.1, domestic ferret XP_004744052.1, rat NP_110502.2, mouse NP_001300926.1, human Origene clone catalog number RG220167. Amino acid positions numbering is according the human 1998 amino acid isoform (NP_001340878.1). (C) SCN1A sequences in donated human brain samples. DNA sequences represent unique (non-PCR duplicate) reads from assay for transposase-accessible chromatin with sequencing (ATAC-seq) from three independent human patient brain samples (H17.26.001, H17.26.003, H18.03.001) profiled in a previous study^31^. (D) Population allele frequencies in human SCN1A via gnomAD database^84,91^. gnomAD v3 and v2 populations represent partially overlapping healthy patient populations subject to genome and/or exome sequencing. The threonine-encoding allele is represented by a T on the + strand (corresponding to A on the - strand) in 73-74% of the population.

**Supplementary Figure 2:**
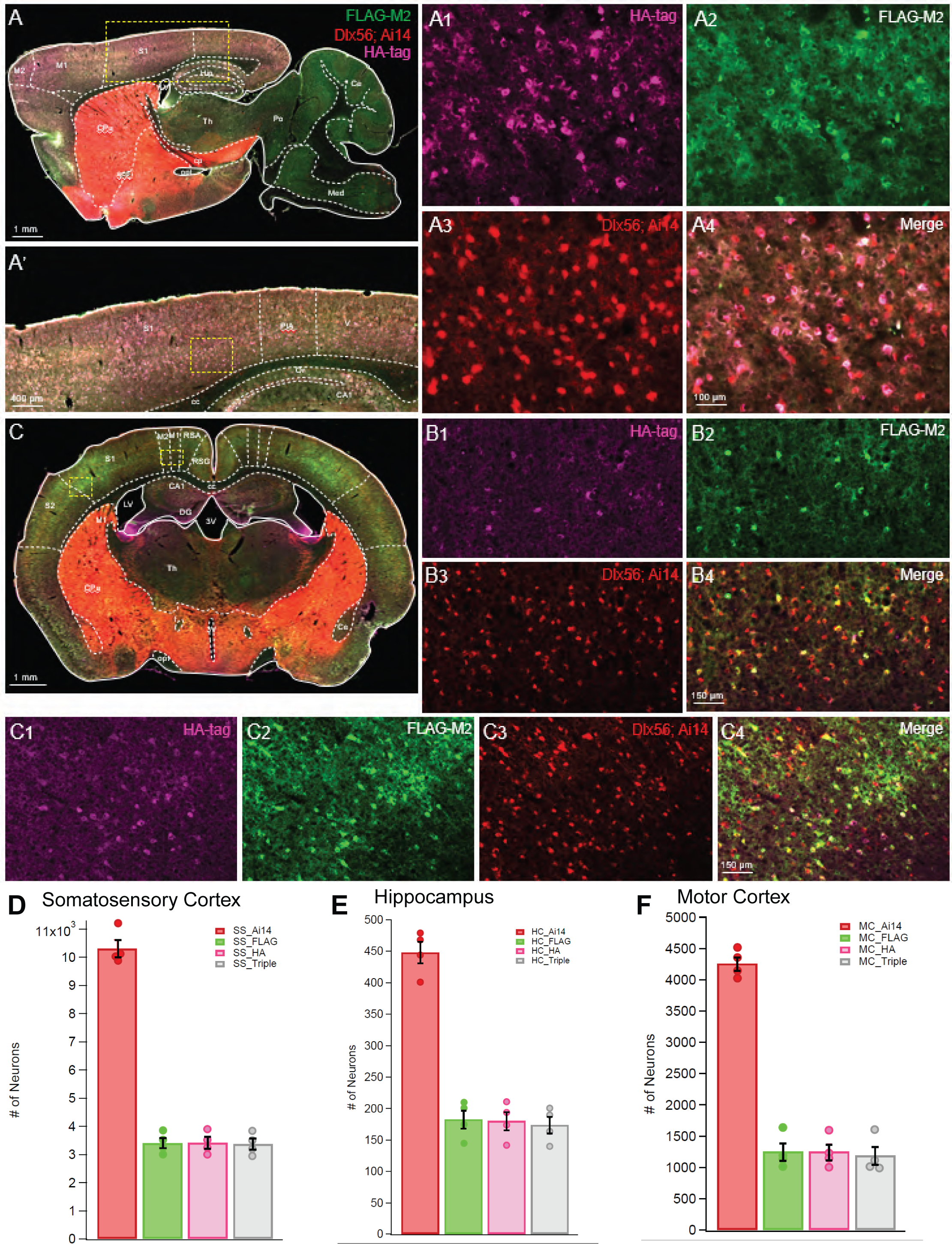
Cell class specificity observed with an independent marker of telencephalic GABAergic interneurons. (A-F) Cell type-specific expression of SCN1A in *Dlx56-Cre; Ai14* reporter mice. We injected these mice at P2 BL-ICV with 3e10 gc each DLX2.0-SCN1A AAV vector produced at PackGene and analyzed expression at P30-P35 with both sagittal and coronal sections (n = 4 mice analyzed, A, C). We analyzed expression specifically in somatosensory cortex (SS, A1-A4 and C1-C4), and motor cortex (MC, B1-B4), and counted absolute numbers of expressing cells in somatosensory cortex (D) and hippocampus (E) and motor cortex (F).

**Supplementary Figure 3:**
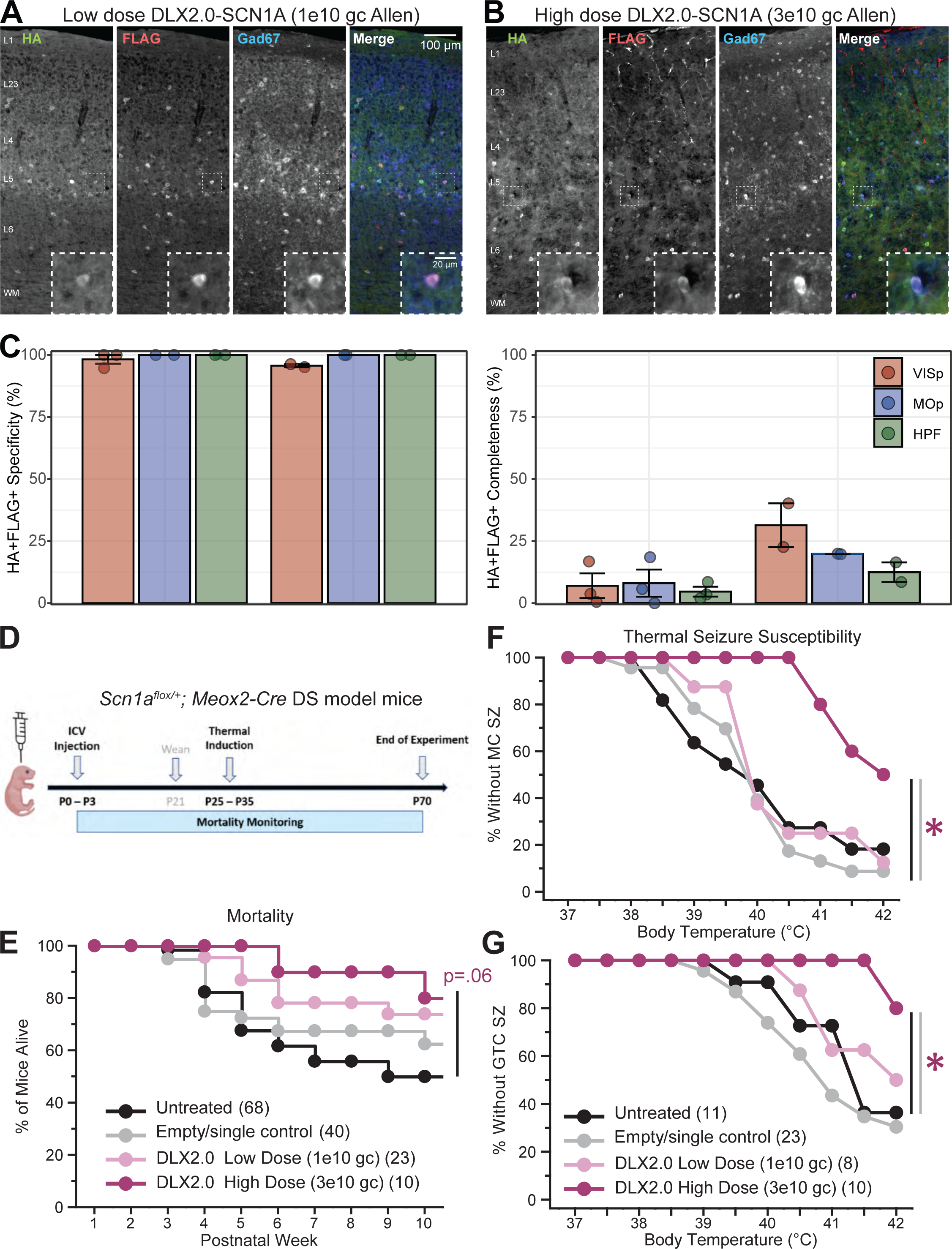
Biodistribution and rescue of epileptic symptoms from independently produced batches of DLX2.0-split intein SCN1A vectors. (A-C) Specific expression of HA-tagged N-terminal and FLAG-tagged C-terminal SCN1A half-channels in Gad67+ neurons with independent packaging of DLX2.0-SCN1A AAVs performed at Allen Institute. Animals were dosed at A) low dose (1e10 gc each vector) or B) high dose (3e10 gc each vector) of DLX2.0-SCN1A AAVs by BL-ICV at P0-P3. C) Telencephalic GABAergic interneuron specificity is maintained while completeness increases with greater dose of AAV. (D-G) Protection from mortality and heat-induced seizures in *Scn1a^fl/+^; Meox2-Cre* mice with independently packaged DLX2.0-SCN1A AAV vectors produced at Allen Institute. (E) Trend towards protection from mortality with high dose of Allen-produced DLX2.0-SCN1A AAV vectors (p = 0.06 by Log-rank test, High dose [n = 10] versus Untreated [n = 68]). (F-G) Dose-dependent protection from heat-induced MC (F) and GTC (G) seizures.* p < 0.05 for comparisons of High dose (n= 10) versus Untreated (n= 11) or Empty/single vector (n= 23) negative controls by Log-rank test. We did not observe significant seizure protection with low doses of Allen Institute-produced DLX2.0-SCN1A AAVs.

**Supplementary Figure 4:**
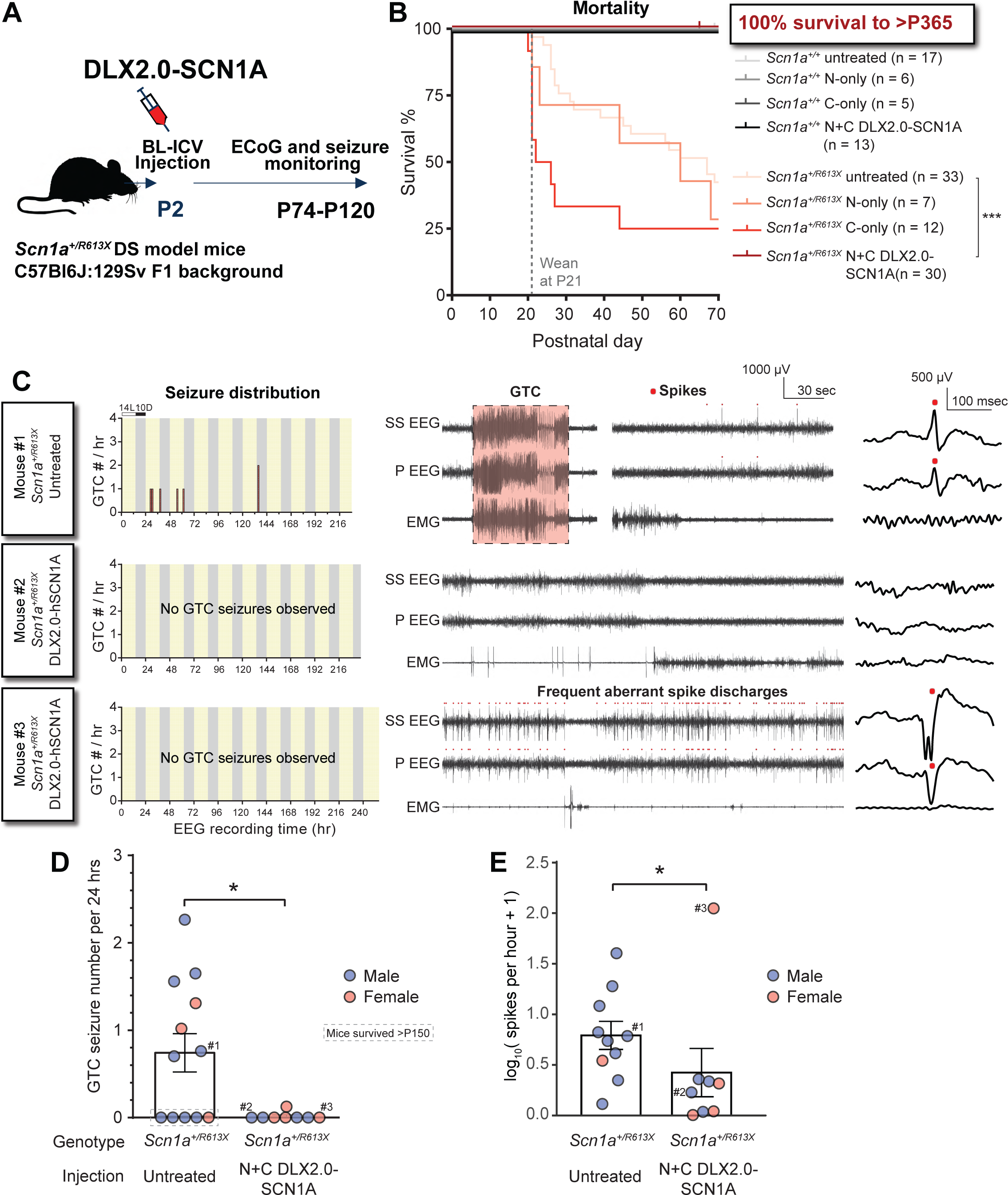
Rescue of mortality and epileptic symptoms in a second independent mouse model of DS. (A) Testing DLX2.0-split-intein-SCN1A in an independent mouse model of DS. We generated *Scn1a^+/R613X^* mice on a mixed F1 C57Bl/6:129Sv background, injected neonates BL-ICV with DLX2.0-split-intein-SCN1A (3e10 gc each vector), and monitored mortality over the first 70 days of life, and then performed ECoG seizure monitoring on rescued animals following mortality monitoring. (B) Mortality monitoring after DLX2.0-split-intein-SCN1A administration. N+C DLX2.0-SCN1A provides highly significant complete rescue from mortality to beyond 365 days. *** p < .001 by Mantel-Cox test (chi-square= 77.69, df= 7) after correction for multiple planned comparisons (3 comparisons of injection materials against untreated *Scn1a^+/R613X^* mice). All other within-genotype comparisons are not significant (p> 0.05) after accounting for multiple comparisons. (C) Seizure monitoring and ECoG in DS model mice. After P70, we implanted headmounts and monitored animals for seizures and epileptic activity by paired ECoG with channels in somatosensory (SS) and parietal (P) cortex, EMG, and video monitoring. Example #1: untreated *Scn1a^+/R613X^* mouse displays several non-uniformly distributed GTC seizures over 228 hours of recording (one GTC seizure shown), as well as interictal spikes marked by red dots (zoom in on one example spike). Example #2 N+C DLX2.0-SCN1A-injected *Scn1a^+/R613X^* mouse displays no GTCs and few spikes. Example #3 N+C DLX2.0-SCN1A-injected *Scn1a^+/R613X^* mouse shows no GTCs but frequent spikes with aberrant spike morphology, likely an outlier. (D) Protection from GTCs with treatment in DS model mice. We manually quantified GTC events over the full recording session for each animal, and displayed results as events per 24 hour recording period, which revealed a significant seizure reduction with administration of N+C DLX2.0-SCN1A. * Mann-Whitney U test p= 0.020; data are non-normally distributed according to Shapiro-Wilk test (untreated p= 0.018; N+C DLX2.0-SCN1A p= 1.0e-6). We also observe that all untreated *Scn1a^+/R613X^* mice with zero GTCs during the recording session eventually survive beyond P150. Marked mice (#1, #2, #3) correspond to the example mice displayed in panel C. (E) Fewer spikes with treatment in DS model mice. We identified spikes over the final day of the recording session for each animal using the line length threshold method^74,82^ (see Methods for details). Quantified spikes are contemporaneous in both SS and P channels, as expected for genetic generalized epilepsy such as DS^19,20,22,24,46,92^. Marked mice (#1, #2, #3) correspond to the example mice in panel C. Despite outlier injected example mouse #3 having aberrant frequent spikes, we observe significantly fewer spikes after injection in the injected mice versus untreated mice (* Mann-Whitney U test p= 0.033). Data are non-normally distributed according to Shapiro-Wilk test (untreated p= 0.0015; N+C DLX2.0-SCN1A p= 1.5e-6).

**Supplementary Figure 5:**
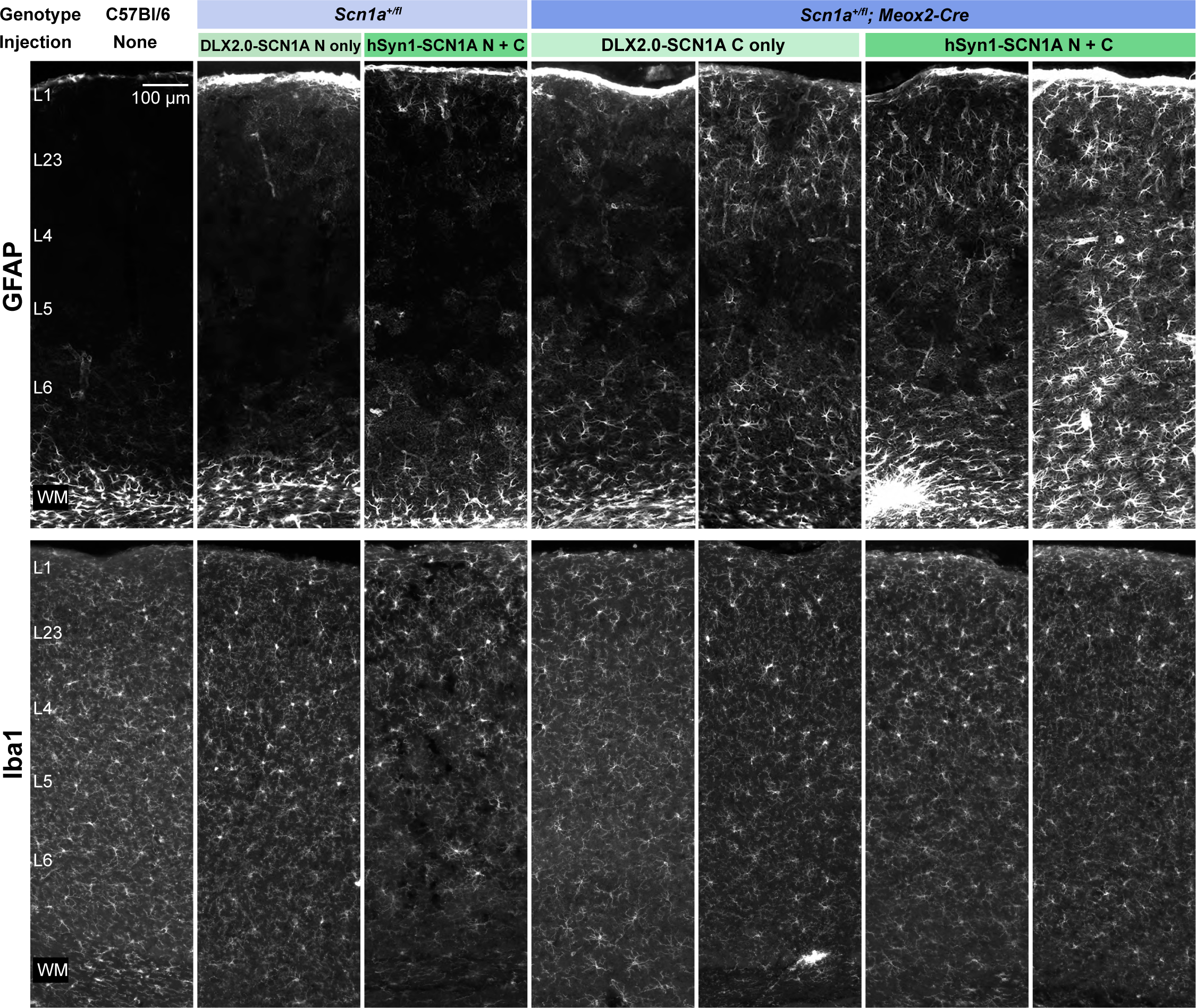
Astrogliosis but not microgliosis induced by nonselective neuronal expression of SCN1A. We sacrificed *Scn1a^+/fl^; Meox2-Cre* DS model and Cre-negative littermate control mice following mortality monitoring after P2 BL-ICV injections (3e10 gc each vector) of hSyn1-driven N+C SCN1A or DLX2.0-driven N-only or C-only single-part negative controls as indicated. Animal ages ranged from P72-P89, and conditions represent two (littermate control) or three (DS model) animals analyzed per condition. We analyzed brains by IHC to assess astrogliosis (GFAP) or microgliosis (Iba1) in cortex (VISp shown). For DS model mice, two example mice spanning the range of astrogliosis observed are shown. Nonselective neuronal expression of SCN1A exacerbated astrogliosis in DS model mice but did not cause overt changes in microglial appearance.

